# Specificities of Chemosensory Receptors in the Human Gut Microbiota

**DOI:** 10.1101/2025.02.11.637667

**Authors:** Wenhao Xu, Ekaterina Jalomo-Khayrova, Vadim M Gumerov, Patricia A. Ross, Tania S. Köbel, Daniel Schindler, Gert Bange, Igor B. Zhulin, Victor Sourjik

**Author notes:** Address correspondence to Victor Sourjik, Igor Zhulin, or Gert Bange. These authors contributed equally to this work (Wenhao Xu, Ekaterina Jalomo-Khayrova).

## Abstract

The human gut is rich in metabolites and harbors a complex microbial community, yet the sensory repertoire of its commensal bacteria remains largely uncharacterized. Here we systematically mapped ligand specificities of extracytoplasmic sensory domains from twenty members of the human gut microbiota, with a primary focus on the abundant and physiologically important class of Clostridia. We identified diverse metabolites as specific stimuli for three major functional classes of transmembrane receptors. We further characterized novel subsets of sensors belonging to the Cache superfamily, specific for lactate, dicarboxylic acids, and for uracil and short-chain fatty acids (SCFAs), respectively, and investigated the evolution of their ligand specificity. Structural and biochemical analysis of the newly described dCache_1UR domain revealed an independent binding of uracil and SCFA at distinct modules. Altogether, we could identify or predict specificities for over a half of the Cache-type chemotactic sensors in the selected gut commensals, with the carboxylic acids representing the largest class of ligands. Among those, the most commonly found specificities were for lactate and formate, indicating particular importance of these metabolites in the human gut microbiome and consistent with their observed beneficial impact on the growth of selected bacterial species.

## Introduction

The gut microbiota, a complex and dynamic assembly of microorganisms residing in the human gastrointestinal (GI) tract, plays a crucial role in host health^1,2^. The composition and resilience of the gut microbiota largely depend on the complex metabolic and signaling interactions of microorganisms with the host and among each other^2,3^, which requires the ability to detect changes in levels of nutrients and signaling molecules^4,5^ using a wide array of signal transduction systems^6^. Major functional families of bacterial environmental sensors are chemotaxis receptors (MCPs) that control bacterial motility in environmental gradients^7,8^, histidine kinases (HKs) that mediate transcriptional responses^9^, and enzymatic sensors including adenylate-, diadenylate- and diguanylate cyclases (DGCs) and phosphodiesterases that regulate the levels of second messengers^10^.

Although these signal transduction pathways have been extensively investigated, beyond a few model organisms little is known about the specificities of transmembrane sensors that bacteria use to perceive their surroundings^6,7,11^. Canonical sensory receptors possess extracytoplasmic ligand-binding domains (LBDs), although various alternative modes of sensory perception have also been described^7,11,12^. LBDs are structurally highly diverse^7^ and function as independent ligand-binding modules that can be shared by different receptor families^13^. Although LBDs can be classified into various families, their ligand specificity is highly variable and evolves rapidly and, in most cases, cannot be predicted solely based on the LBD sequence or structure^7,14^. The reported repertoire of known signals perceived by sensory receptors across bacterial species is remarkably broad and organism-specific, including various nutritionally valuable metabolites, bacterial and eukaryotic signaling molecules, as well as ions, gases, pH, light, and various cellular stresses^15,16^. These sensory capabilities are known to be intimately associated with their living environment^17,18^, but which of the numerous host-derived and bacteria-derived metabolites^19,20^ serve as preferred signals for the gut bacteria remains unknown.

To systematically identify these signals, we selected around one hundred representative LBDs from the previously reported dataset of extracytoplasmic sensory domains derived from the human gut microbiome^21^, of which seventy proved amendable to the subsequent biochemical analysis. We then investigated their binding specificities for a set of over 150 metabolites that are known to be present in the mammalian gut. We focused on bacteria from the class of Clostridia due to their abundance in the gut microbiota and their critical role in the maintenance of gut homeostasis^22^. Additionally, this class contains the most prevalent chemotactic bacterial species in the gut microbiota, and thus covers all three major functional families of bacterial environmental sensors. We identified 34 interactions of LBDs with various ligands, including short-chain fatty acids (SCFAs), lactate, formate, indole, pyrimidines, amines, purines, and amino acids. The ability of selected ligands to elicit signaling responses was confirmed by constructing chimeric receptors that could be studied in the heterologous *Escherichia coli* system. Structural analysis and mutagenesis were used to characterize ligand-binding sites for the two prominent groups of domains with novel specificities, enabling insights into their evolution and rational modification of ligand specificities. Most identified interactions appear to reflect chemotactic responses to metabolically relevant compounds that impact bacterial growth, highlighting the physiological importance of chemotaxis toward nutrients in the gut microbiota.

## Results

### Construction of the LBD library and high-throughput ligand screening

We selected an LBD library including over one hundred extracytoplasmic sensory domains from 20 different human gut bacteria, using our previously established dataset for the human gut microbiome^21^ (Supplementary Table 1 and Supplementary Data 1). The majority of the selected organisms belong to Clostridia, a highly represented and ecologically significant class of bacteria in the human gut microbiome, which includes the most abundant motile commensal bacteria. These species thus harbor all three major families of bacterial environmental sensors, with the majority of the LBDs in our library being derived from MCPs closely followed by HKs, and with DGCs representing the smallest proportion (Extended Data Fig. 1a). Our LBD library consists of three structural groups of domains, including dCache (dual or bimodular) and sCache (single or monomodular) domains of the Cache superfamily and 4HB (four-helix bundle) domains (Extended Data Fig.1b), which is similar to the distribution of these groups within the initial dataset^21^. Also consistent with that dataset, the distribution of domain groups differs between the receptor families, with 4HB domains more frequent in MCPs and Cache domains being more frequent in HKs (Extended Data Fig. 1c-d).

We next expressed and purified LBD proteins from the library and subjected them to thermal shift assays (TSA) for high-throughput ligand screening^23^. Most purified LBDs exhibited distinct melting curves as the temperature increased (Extended Data Fig. 1e), confirming that these proteins were initially well-folded. Their specificities could therefore be characterized based on the changes in thermal stability upon ligand binding. The resulting set of 70 purified and folded LBDs had similar representation of the receptor families as the original library (Fig. 1a, Supplementary Data 1, and Extended Data Fig.1a, f-g), despite some enrichment of the Cache domains (Fig. 1b and Extended Data Fig.1b).

**Fig. 1.**
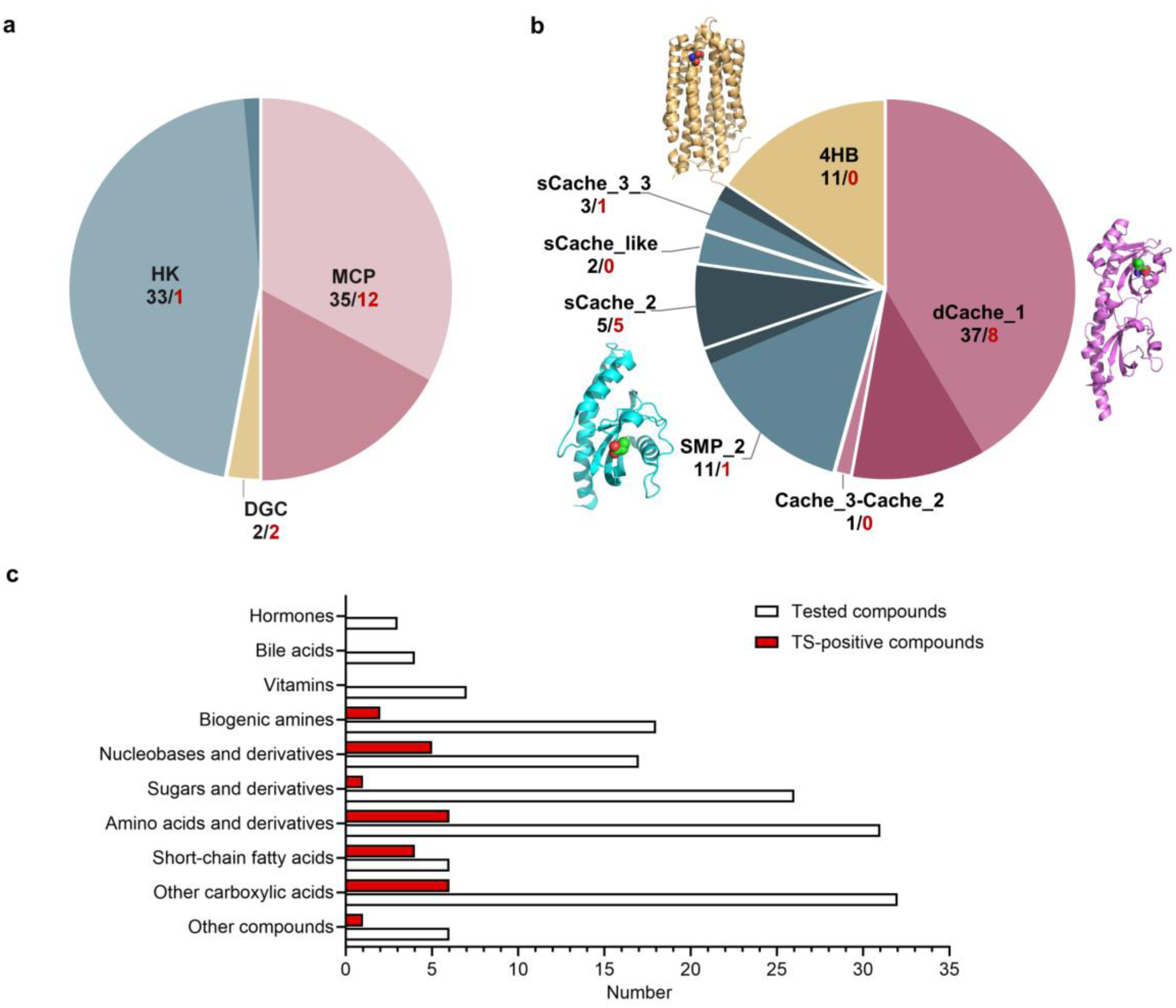
Overview of the high-throughput ligand screening for the LBD and ligand libraries. **a**, **b**, Composition of the folded LBDs presented by the receptor type (a) and by the domain family (b), respectively. The numbers below receptor and domain names indicate studied LBDs (black) and those with identified ligands (red). Double-module Cache domain family (dCache) is depicted in pink, single-module Cache domain family (sCache) is represented in blue, and the four-helix bundle_MCP family (4HB) is illustrated in yellow. Fractions of LBDs with identified ligands within different groups are highlighted in darker color. Domain definitions follow the Pfam domain nomenclature. MCP, methyl-accepting chemotaxis protein; HK, histidine kinase; DGC, diguanylate cyclase. Three-dimensional structures of three structural domain families are shown in Fig. 1b: 4HB – Tar bound to L-aspartate (PDB ID 4Z9H), dCache – PctA bound to L-isoleucine (PDB ID 5T65), and sCache – PscD bound to propionate (PDB ID 5G4Z). **c**, Composition of all studied compounds from two human gut metabolite (HGMT) plates and compounds identified as ligands of one or multiple LBDs using thermal shift assays (TSA; referred to as TS-positive compounds). Layouts of two HGMT plates are shown in Supplementary Table 2.

To screen ligands for these LBDs, we used the custom-assembled human gut metabolite (HGMT) plates containing over 150 different chemical compounds, mainly nutrients but also several animal hormones and other compounds known to be present in the human gut (Fig. 1c and Supplementary Table 2). We observed the binding of one or several compounds to multiple LBDs, including most prominently SCFAs and other carboxylic acids, as well as amino acids and nucleobases and their derivatives (Fig. 1c, Extended Data Fig. 2a-b, and Table 1). Interestingly, while we successfully identified ligands for a large fraction of the tested dCache and sCache domains, none of the proteins in the 4HB group exhibited thermal shifts upon the addition of compounds (Fig. 1b). Nevertheless, our TSA setup can be used to characterize ligand binding to the 4HB LBDs, as confirmed for 4HB LBDs of *E. coli* chemoreceptors Tar and Tsr (Extended Data Fig. 2c-d).

**Table 1.**
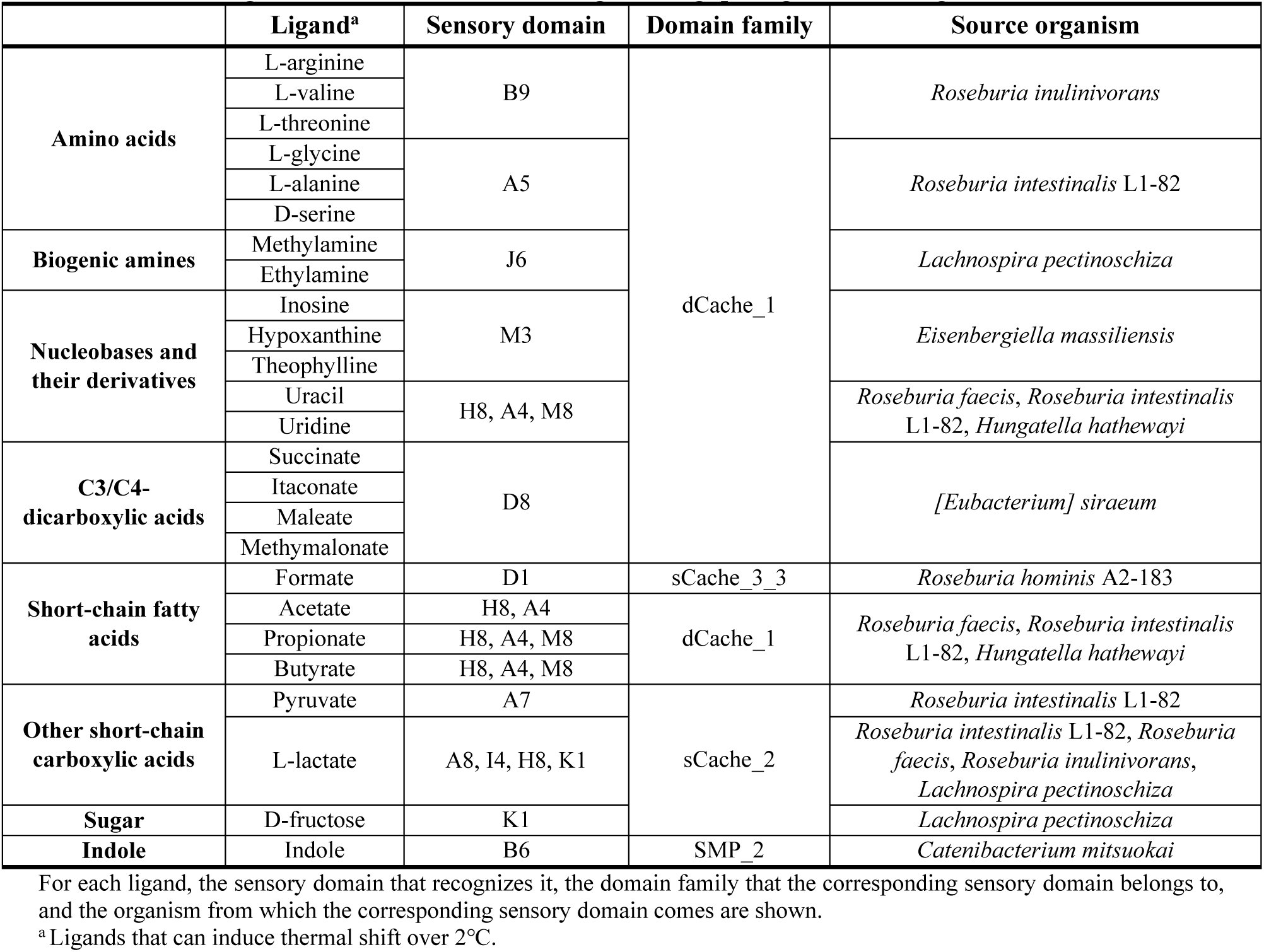
Putative ligands identified in the initial high-throughput ligand screening via TSA.

Altogether, we could assign ligands for 15 LBDs (Table 1). Interestingly, despite having a similar number of well-folded LBDs, HKs and MCPs showed a striking difference in their ligand-binding properties. Whereas 12 MCP LBDs could bind diverse chemical compounds from the HGMT plate, only one HK LBD could be assigned a ligand, indole (Fig. 1a). In addition, nucleobases and their derivatives could be assigned as ligands to both tested DGC LBDs.

### Lactate is a prevalent ligand for sCache_2 domains in the human gut microbiota

We assigned several compounds as specific ligands for seven sCache-type LBDs, including L-lactate or pyruvate for five sCache_2 MCP domains, formate for an sCache_3_3 MCP domain which contains the recently reported formate binding residues^24^, and indole for the SMP_2 HK domain (Fig. 2a and Table 1). The binding of L-lactate to the K1 and C1 LBDs (Fig. 2b) and the binding of indole to the B6 LBD were confirmed by isothermal titration calorimetry (ITC) measurements (Fig. 2c). The ability of the K1 LBD to mediate chemotactic signaling upon L-lactate binding was further confirmed using the K1-Tar chimera, where this LBD was fused to the signaling domain of Tar. When expressed in the chemoreceptor-less *E. coli* strain, K1-Tar indeed mediated chemotactic responses to L-lactate, as determined using the pathway activity assay based on the Förster resonance energy transfer (FRET)^25,26^ (Fig. 2d). No response was observed for the same background strain expressing the native Tar receptor (Supplementary Fig. 1a), confirming its specificity.

**Fig. 2.**
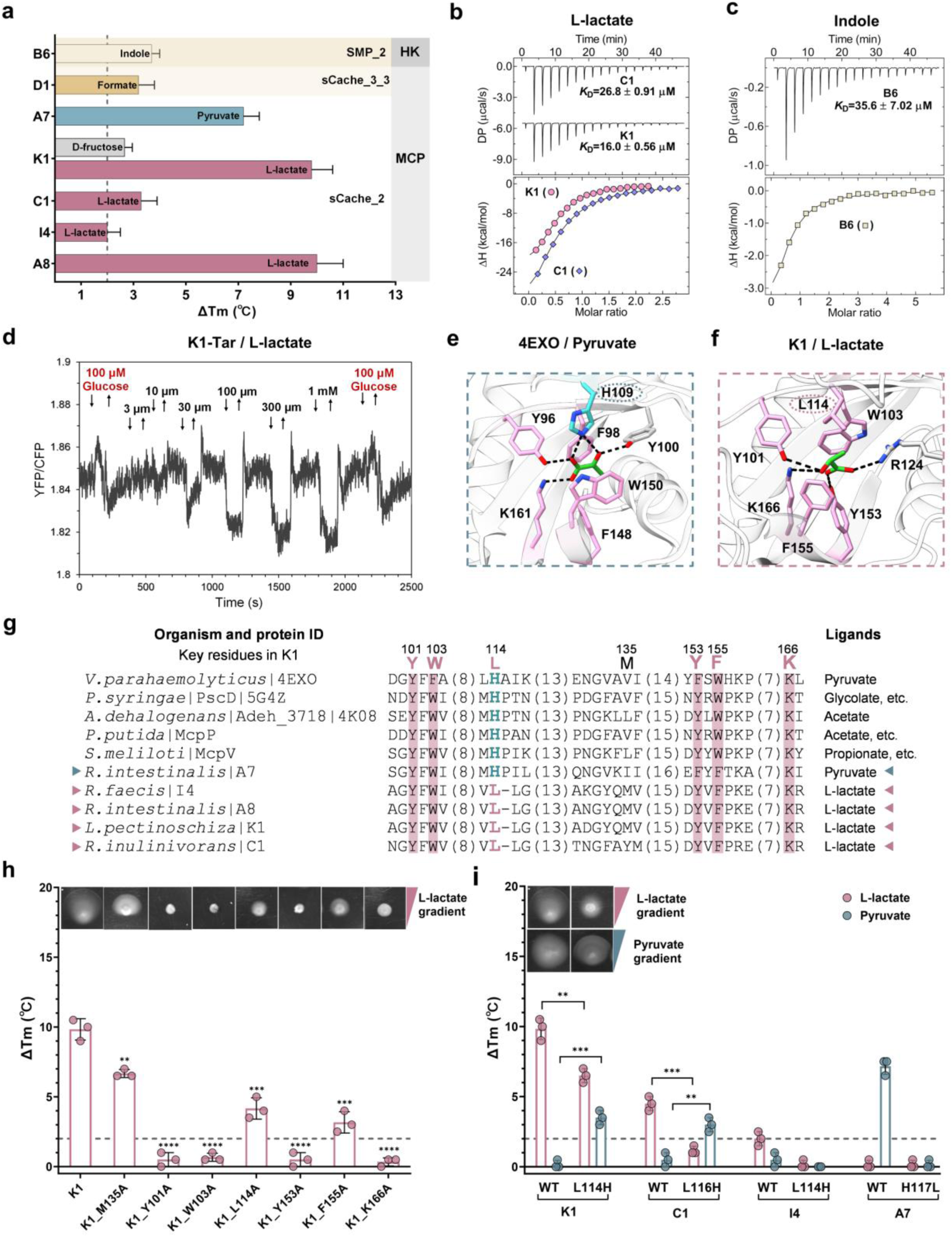
Characterization of the sCache_2 domains specific for short-chain carboxylic acids. **a**, Thermal shifts observed for sCache domains upon exposure to 2 mM final concentrations of respective ligands. Subfamilies of domains and types of receptors are indicated. The gray dashed line represents the 2°C threshold used as the significance cutoff for ligand identification. Data are presented as the mean ± standard deviation from three independent biological replicates. **b**, **c**, ITC measurements of the K1 and C1 LBDs binding to L-lactate (b) and of the B6 LBD binding to indole (c). Upper panels: Raw titration data. Lower panels: Integrated, dilution heat corrected, and concentration normalized raw data. The lines are the best fits using the “One binding site” model, with the derived dissociation constants (*K*_D_) being indicated. Further experimental details are provided in Supplementary Table 5. **d**, FRET measurements of the K1-Tar hybrid chemoreceptor response to L-lactate. The buffer-adapted *E. coli* cells expressing the CheZ-CFP/CheY-YFP FRET pair and K1-Tar as the sole receptor were stimulated by stepwise addition (down arrow) and subsequent removal (up arrow) of the indicated concentrations of L-lactate. D-glucose was used in FRET measurements as a positive control to confirm the activity of the K1-Tar chimera, as it stimulates the cytoplasmic domain of the chemotaxis receptors. **e**, Reported co-crystal structure of the sCache_2 domain in complex with pyruvate (PDB ID 4EXO). **f**, Computational docking of L-lactate to the AlphaFold3 model of K1 LBD. The conformation with the highest confidence is presented. Residues involved in ligand binding are shown in stick mode. The key residues shown in pink indicate conserved amino acids, while variable residues are circled with dashed lines, differentiated by color: blue for H109 in 4EXO and pink for L114 in K1. Black dashed lines present hydrogen bonds. **g**, Protein sequence alignment of experimentally studied sCache_2 domains that bind short-chain carboxylic acids (SCCAs), with the conserved residues highlighted in pink. Newly characterized sensory domains are marked with triangles. The numbers above the alignment correspond to the amino acid positions in K1. **h**, Binding and chemotaxis measurements of K1 LBD and its indicated alanine-substituted mutants in response to L-lactate. M135A substitution was used as a negative control. **i**, Binding and chemotaxis measurements of the indicated LBDs. The chemotaxis measurements in (h, i) were performed on soft-agar plates with gradients of either L-lactate or pyruvate, indicated by pink or blue triangles. Thermal shifts induced by 2 mM L-lactate are shown in pink, and those by 2 mM pyruvate are shown in blue. The gray dashed line indicates the threshold of 2°C used as a significance cutoff for ligand identification. The data represent the mean ± standard deviations of three independent biological replicates. Each data point corresponds to an independent biological measurement. Asterisks denote statistically significant differences (unpaired *t*-test): **p ≤ 0.01, ***p ≤ 0.001, and ****p ≤ 0.0001.

Given that all characterized sCache_2 MCP domains exhibited specificity toward L-lactate or pyruvate, there is apparently a subset of sCache_2 domains in the human gut microbiota dedicated to recognizing C2/3 carboxylic acids. We identified the structure of a K1-homologous sCache_2 domain from *Vibrio parahaemolyticus* bound to pyruvate (PDB ID 4EXO) in the RCSB protein data bank^27^ (Fig. 2e). Comparison with the molecular docking results for L-lactate binding to the AlphaFold3 model of K1 LBD (Fig. 2f) revealed that both ligands share the same binding mode to their LBDs, with two aromatic amino acids (4EXO^F98^/K1^W103^ and 4EXO^W150^/K1^F155^) sandwiching the ligand, and three residues (4EXO^Y96^/K1^Y101^, 4EXO^F148^/K1^Y153^, and 4EXO^K161^/K1^K166^) forming hydrogen bonds with carboxyl oxygen atoms of the ligand. Indeed, these residues are highly conserved across all sCache_2 domains that were shown to bind short-chain carboxylic acids, either in the literature or in our screen (Fig. 2g). However, the histidine residue (H109) that also establishes a direct contact with pyruvate in 4EXO, as well as in two other sCache_2 domains, PscD and Adeh_3718, where ligand binding modes have been established^28,29^, is replaced in all identified L-lactate sensors by leucine (L114 in K1; Fig. 2g), which does not directly contribute to L-lactate binding in the structural model of K1 LBD (Fig. 2f).

To verify the role of the corresponding amino acid residues in L-lactate binding, we replaced them individually with alanine. Substitutions at any of the four highly conserved residues (Y101, W103, Y153, and K166; numbered according to K1) completely abolished L-lactate binding as measured by TSA, while substitutions at F155 and L114 strongly reduced binding (Fig. 2h). A lesser effect was observed for the substitution at M135 that is not conserved among these sensory domains. These *in vitro* binding results were validated by chemotaxis assays for the corresponding alanine-substituted mutants of the K1-Tar chimeric receptor using soft-agar gradient plates^26^ (Fig. 2h *Inset*)

Since the presence of L114 instead of histidine within the binding pocket appears to correlate with the binding of L-lactate, we hypothesized that this difference may be important for ligand selectivity. Indeed, the replacement of leucine with histidine at this position in the K1 and C1 LBDs enabled binding of pyruvate but weakened binding of L-lactate (Fig. 2i and Supplementary Fig. 2). This observation was further confirmed by the chemotaxis assay for the K1-Tar_L114H mutant (Fig. 2i *Inset*). Thus, we identified L114 in the ligand-binding pocket of sCache_2 domains as the specificity determinant for L-lactate binding.

Although L-lactate, primarily produced by host epithelial cells, is the predominant lactate enantiomer in the GI tract, gut bacteria can generate both L- and D-lactate^30^. We thus also examined the binding of D-lactate to the K1 LBD. D-lactate elicited a significant but weaker thermal shift in the K1 LBD compared to L-lactate (Extended Data Fig. 3a) and only a weak response in the FRET assay (Extended Data Fig. 3b). Consistently, K1 LBD exhibited a fourfold lower binding affinity for D-lactate than for L-lactate (Extended Data Fig. 3c and Fig. 2b), suggesting that L-lactate is the preferred ligand of this subset of sCache_2 domains.

### Diverse ligands bind to the membrane-distal modules of dCache_1 domains

The spectrum of ligands identified for dCache_1 domains exhibited greater diversity, including amino acids, purines, pyrimidines, biogenic amines, and carboxylic acids (Fig. 3a and Table 1). The membrane-distal modules of MCP amino acid sensors A5 and B9, the MCP amine sensor J6, and the DGC purine sensor M3 were found to possess specificity motifs within their putative pockets, as recently characterized for dCache_1 domains^31–33^ (Extended Data Fig. 4). Thus, these ligands likely bind to the membrane-distal modules of sensory domains. The binding of L-threonine to the B9 LBD, as well as ethylamine and methylamine to the J6 LBD, was confirmed by ITC (Fig. 3b-c). Furthermore, the J6-Tar chimeric receptor mediated specific responses to methylamine and ethylamine when measured by FRET in *E. coli* (Fig. 3d and Supplementary Fig. 1b), in contrast to the native Tar (Supplementary Fig. c-d).

**Fig. 3.**
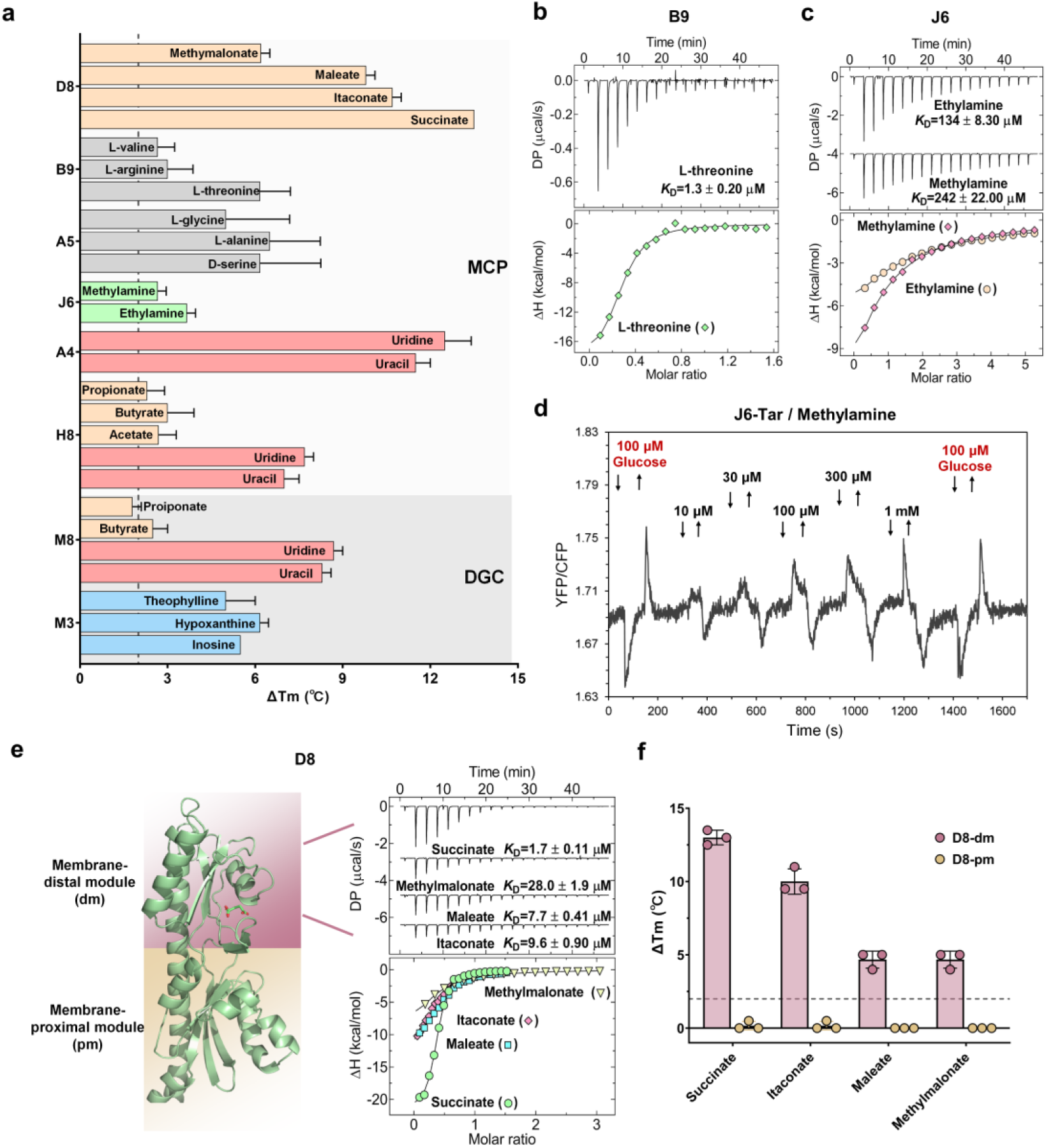
Characterization of diverse ligands binding to the membrane-distal module of dCache_1 domains. **a**, Thermal shift observed for dCache_1 domains upon exposure to 2 mM concentrations of respective ligands, with subfamilies of domains and types of receptors being indicated. The gray dashed line represents the 2°C threshold used as the significance cutoff for ligand identification. Data are presented as the mean ± standard deviation from three independent biological replicates. **b**, **c**, ITC measurements for the B9 (c) and J6 (d) LBD binding to indicated ligands. Upper panels: Raw titration data. Lower panels: Integrated, dilution heat corrected, and concentration normalized raw data. The lines are the best fits using the “One binding site” model, with the derived dissociation constants (*K*_D_) being indicated. **d**, FRET measurements of the J6-Tar hybrid chemoreceptor response to methylamine. The buffer-adapted *E. coli* cells expressing the CheZ-CFP/CheY-YFP FRET pair and J6-Tar as the sole receptor were stimulated by stepwise addition (down arrow) and subsequent removal (up arrow) of the indicated concentrations of methylamine. **e**, ITC measurements of the D8 LBD binding to indicated ligands, with the derived dissociation constants (*K*_D_) being indicated. Further experimental details are provided in Supplementary Table 5. The AlphaFold3 model of the D8 LBD in complex with docked succinate is shown on the left, with the membrane-distal module (dm) colored pink and the membrane-proximal module (pm) colored yellow. **f**, Thermal shift measurements for the membrane-distal module of the D8 LBD (D8-dm) and the membrane-proximal module of the D8 LBD (D8-pm) in the presence of 2 mM concentrations of indicated compounds. The gray dashed line indicates the threshold of 2°C used as a significance cutoff for ligand identification. The data represent the means ± standard deviations of three biologically independent replicates. Each data point corresponds to an independent biological measurement.

Thermal shift assays demonstrated the binding of the D8 LBD to four C3/C4 dicarboxylic acids (Fig. 3a), which was further confirmed by ITC (Fig. 3e). Despite their general metabolic importance for bacteria, no Cache-superfamily receptors specific for these dicarboxylic acids have been previously characterized. Thus, to identify which module of the D8 LBD binds these ligands, we separately tested the membrane-distal module (D8-dm) and the membrane-proximal module (D8-pm). Thermal stabilization by all four ligands was observed for the membrane-distal module but not for the membrane-proximal module (Fig. 3f), suggesting that binding occurs in the pocket of the membrane-distal module in this case as well.

### Uracil and SCFAs independently bind to different modules of dCache_1UR

Another novel subset of dCache_1 domains, represented by the H8 and A4 MCP sensors and by the M8 DGC sensor, was observed to bind two distinct classes of compounds: (1) uracil and its derivative uridine, which exhibited larger thermal shifts and higher binding affinities, and (2) SCFAs, which showed smaller thermal shifts and lower binding affinities (Fig. 3a and Extended Data Fig. 5a-e). We therefore named this subset the dCache_1UR domains. The H8 LBD was also able to bind another uracil derivative, the anticancer agent 5-fluorouracil^34^ (Extended Data Fig. 5b). Given the difference between molecular structures of SCFAs and uracil, we reasoned that these ligands are likely to bind at different sites. Indeed, the membrane-distal modules of dCache_1UR domains could bind uracil but not SCFAs (Extended Data Fig. 5f-g), indicating that the binding of SCFAs might instead occur at the membrane-proximal module.

To elucidate the molecular basis of ligand recognition, we crystallized the purified A4 LBD in the presence of uracil and propionate. The resulting structure was determined at a resolution of 1.46 Å (Fig. 4a-c). The A4 domain comprises two modules: (1) the membrane-distal module (residues 57-206) containing seven β-strands, four α-helices, and half of the α1 helix that connects these two modules, and (2) the membrane-proximal module (residues 35-56, 207-303) formed by four β-strands, five α-helices and the other half of the aforementioned α1 helix (Extended Data Fig. 6a). This organization is similar to that of other dCache_1 domains, although structural details of these domains can vary^14,15^. Within the membrane-distal module, an electron density that could be unambiguously assigned to uracil was observed (Extended Data Fig. 6b). Uracil is tightly coordinated by eight key residues and one water molecule (Fig. 4b and Extended Data Fig. 6b). The residues R116, T145, N178, and N180 stabilize uracil via hydrogen bonds at a distance of ∼ 2.9 Å. Additionally, a water molecule forms hydrogen bonds in a tetrahedral coordination with uracil and residues Y176, N178, and D205, with respective distances of 2.8 Å, 3.1 Å, 2.9 Å, and 2.7 Å (Fig. 4b). Moreover, F129 and W160 form π-π stacking interactions with uracil at distances of 3.8 Å and 3.6 Å (Fig. 4b). In the proximal module, although propionate was added during crystallization and acetate was absent in the experimental conditions, an acetate molecule was unambiguously explaining the obtained extra density (Extended Data Fig. 6c). Therefore, this acetate molecule was likely endogenous and prebound to the A4 domain during its expression in *E. coli*. Four residues coordinate acetate via hydrogen bonds: Y225, H238, Y273, and K280, with bond distances of 2.6 Å, 2.7 Å, 2.5 Å and, 2.9 Å, respectively (Fig. 4c). Consistently, the AlphaFold3 models of the M8 and H8 LBDs closely resemble the structure of the A4 LBD, with key residues highly conserved (Extended Data Fig. 5h-l). Further analysis of the purified A4 LBD also revealed the presence of endogenously bound uracil (Extended Data Fig. 7a), suggesting that both binding pockets of A4 are at least partially occupied. To eliminate the likely effects of partial occupancy on binding measurements, prebound ligands were removed from the purified protein to obtain a ligand-free A4 LBD (apo-A4; Extended Data Fig. 7a). Apo-A4 LBD indeed exhibited a much higher affinity (Fig. 4d and Extended Data Fig. 5a) and greater thermal shifts to uracil compared to the partially occupied A4 LBD (Extended Data Fig. 7b-c).

**Fig. 4.**
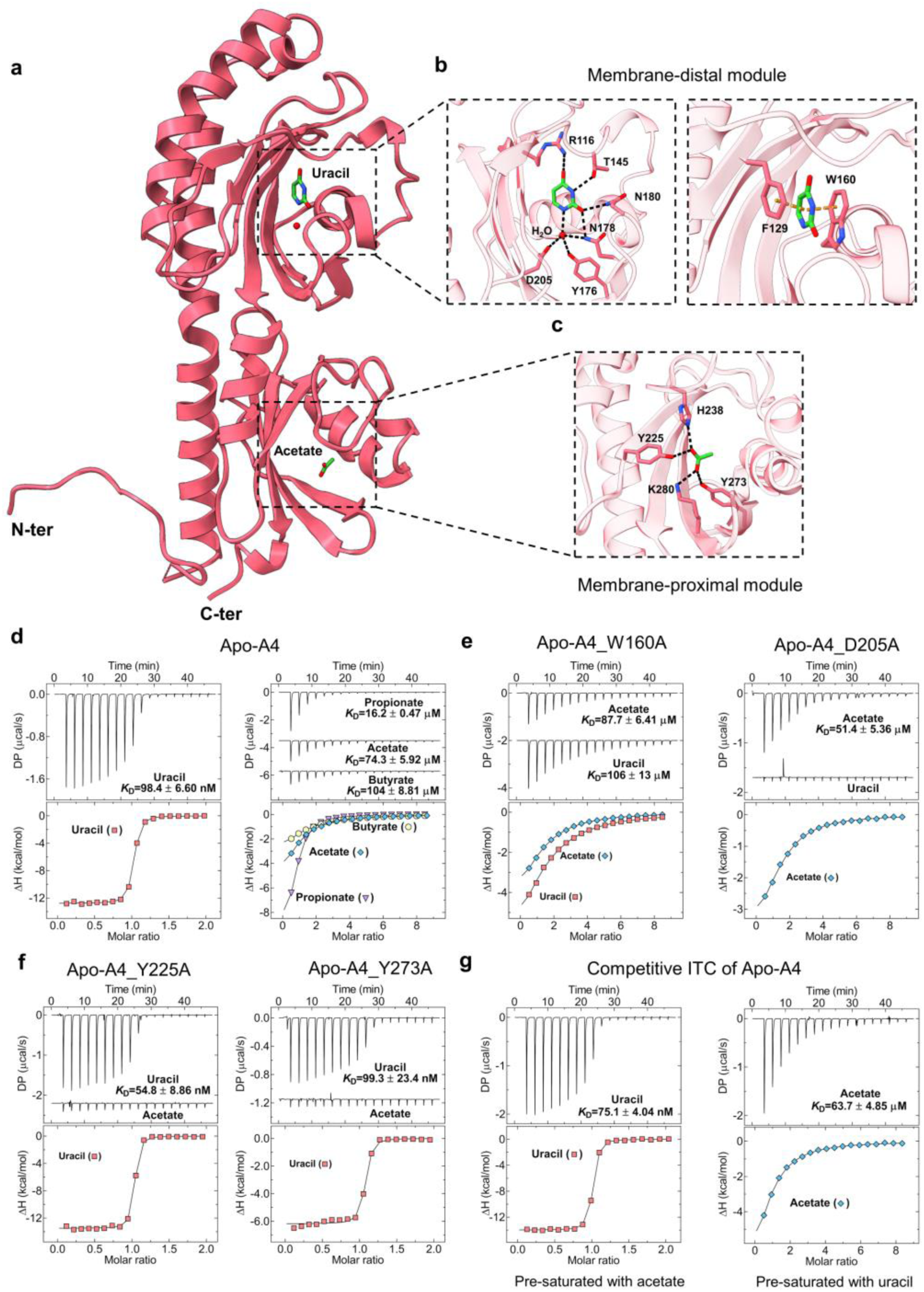
dCache_1UR domain independently binds uracil and SCFAs at distinct modules. **a**, Overall structure of the dCache_1UR domain A4 LBD in complex with uracil and acetate (PDB ID 9HVJ). **b**, **c**, Molecular details of uracil recognition sites at the membrane-distal module (b) by hydrogen bonds (black dashed lines, left) and π-π stacking interactions (orange dashed lines, right), and acetate recognition site at the membrane-proximal module (c) by hydrogen bonds (black dash lines). Residues involved in ligand binding are shown in stick mode. **d**, ITC measurements of Apo-A4 LBD binding to uracil and SCFAs**. e**, **f**, ITC measurements for indicated variants of Apo-A4 with amino acid substitutions in the membrane-distal (e) or membrane-proximal module (f) with indicated ligands. **g**, Competitive ITC measurements of uracil binding to Apo-A4 pre-saturated with acetate (left) and of acetate binding to Apo-A4 pre-saturated with uracil (right). Upper panels: Raw titration data. Lower panels: Integrated, dilution heat corrected, and concentration normalized raw data. Continue lines correspond to the best fit using the “One binding site” model, with the derived dissociation constants (*K*_D_) being indicated. Further experimental details are provided in Method and Supplementary Table 5.

To investigate the contribution of individual amino acid residues in the pocket to uracil binding, we tested the effects of alanine substitutions at these residues. The replacement of R116, F129, W160, and D205 residues resulted in a largely reduced thermal shift (Extended Data Fig. 7b-c) and a strongly reduced binding affinity, or even complete abolishment of binding (Fig. 4e), whereas replacement of other four residues had a more modest impact. Similarly, alanine substitution mutants of four key residues in the SCFA binding pocket disrupted binding (Fig. 4f and Extended Data Fig. 7b-c).

Next, we further investigated whether an allosteric regulation might exist between the binding of SCFA and uracil to the dCache_1UR domain, using competitive ITC (see Methods). No apparent impact of the presence of acetate was observed on the binding of uracil to the A4 LBD or vice versa, suggesting that these ligands bind independently (Fig. 4g). Consistent with that, introducing single point mutations in the binding pocket for one ligand had no impact on binding of the other ligand (Fig. 4e-f and Extended Data Fig. 7b-c).

### Evolutionary relatedness between dCache_1UR and other characterized dCache_1 domains

To explore the evolutionary origin of uracil sensors, we first performed the sequence alignment for newly identified dCache_1UR domains (A4, H8, and M8) with the previously characterized dCache_1AM (McpX), dCache_1AA (PctA), and dCache_1PU (McpH) domains (Fig. 5a). It suggested that dCache_1UR domains shared highest similarity to dCache_1AM domains. To further elucidate the phylogenetic relationships between the dCache_1UR domain and other three dCache_1 domains, we analyzed corresponding protein sequences representing diverse taxonomic groups with variations in their motifs (Supplementary Data 3). Phylogenetic inference (Fig. 5b and Supplementary Fig. 3), performed using a Bayesian approach (see Methods), suggested that the dCache_1AA domain represents the common ancestor of the other three subsets of dCache_1domains. The dCache_1UR and dCache_1AM domain families form two closely related clades that likely evolved from a shared ancestor. In contrast, dCache_1PU domains form a separate, well-defined branch. This is consistent with the previous analyses, which indicated that the dCache_1AA domain represents an ancient, broadly distributed domain family^33^, with amino acid substitutions and structural adaptations in its distal pocket likely driving the diversification and proliferation of dCache_1 domain families with different ligand specificities.

**Fig. 5.**
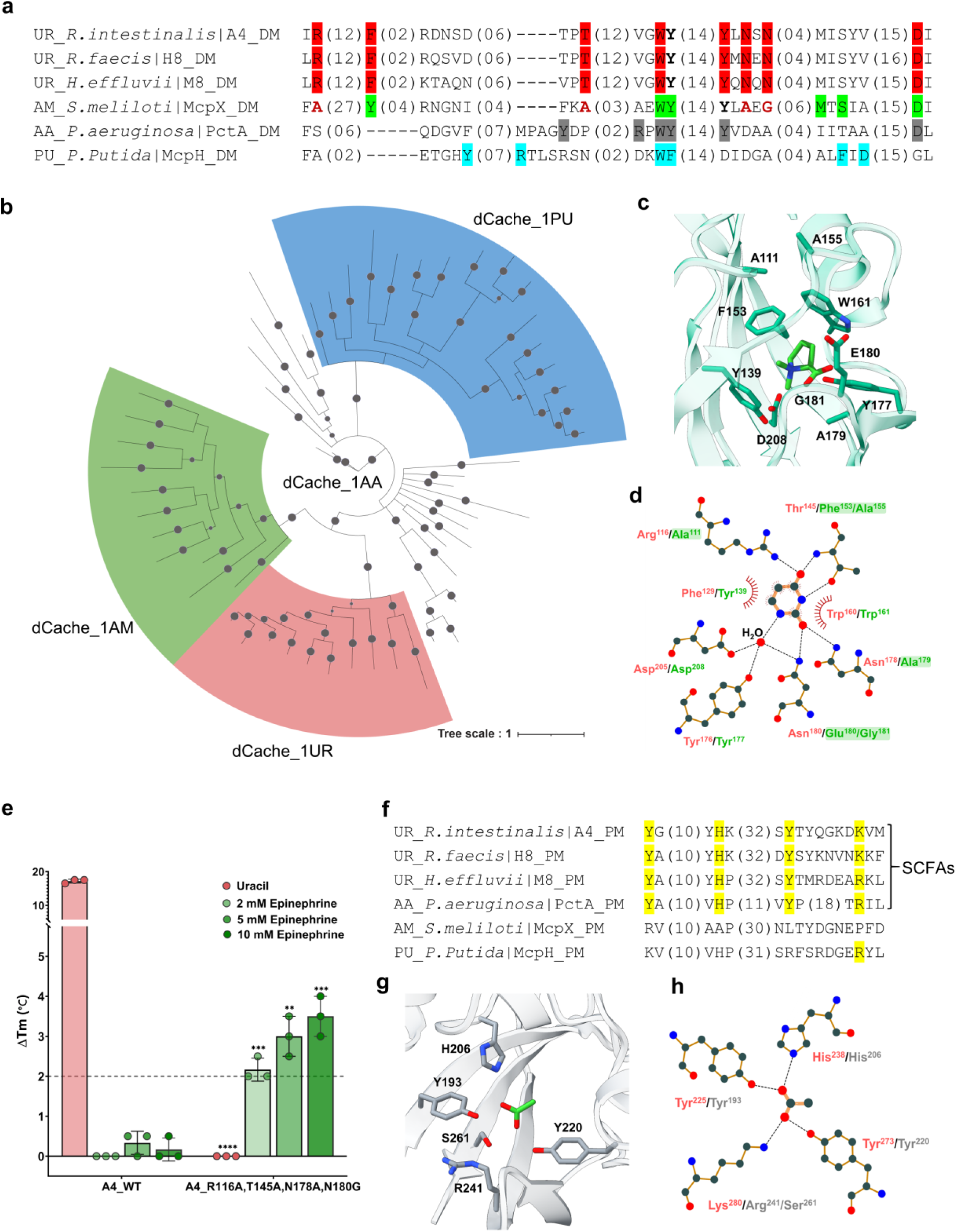
Evolutionary relationships and ligand-binding specificity of dCache_1 domains. **a**, Multiple sequence alignment of the membrane-distal module of uracil, amine, amino acid, and purine sensors. The corresponding motifs are shown in red (uracil-binding key residues), green (amine-binding key residues), gray (amino acid-binding key residues), and blue (purine-binding key residues). The four residues of McpX, shown in bold red, are in positions corresponding to the uracil-binding key residues. **b**, Phylogenetic analysis of the indicated dCache_1 domains. Uracil sensors are marked in red, amine sensors in green, and purine sensors in blue; amino acid sensors are not colored. Probabilities greater than 0.8 are represented by filled gray circles. dCache_1 amino acid sequences from major phyla of each shown family were used. A detailed phylogenetic tree, including information about the proteins used and their corresponding phyla, is provided in Supplementary Data 3. **c**, Amine binding pocket of McpX (PDB ID 6D8V) in complex with proline betaine (PBE). **d**, Schematic representation of residues involved in uracil binding. The residues coordinating uracil in A4 are shown in red, while the corresponding residues in McpX are depicted in green. Residues required for uracil binding but not conserved in McpX are highlighted with a green shadow. Dashed lines represent hydrogen bonds, and half-circles denote hydrophobic interactions. This figure was generated using LigPlot^+^. **e**, Thermal shifts observed for the A4 LBD and A4 _R116A, T145A, N178A, N180G LBD mutants exposed to indicated concentrations of uracil or epinephrine. The data represent the means ± standard deviations of three biologically independent replicates. Each data point corresponds to an independent biological measurement. **f**, Multiple sequence alignment of the membrane-proximal modules from uracil, amine, amino acid, and purine sensors. Amino acids involved in SCFA binding are shown in purple. **g**, Acetate binding pocket of the amino acid sensor PctA (PDB ID 5T65) in complex with acetate. **h**, Schematic representation of residues involved in acetate binding. The residues that coordinate the acetate are shown in red (A4) and gray (PctA). Dashed lines represent hydrogen bonds. This figure was generated using LigPlot^+^.

Consistent with their close relatedness, structural superposition of the ligand binding pockets of McpX and A4 showed that four conserved residues (A4^F129^/McpX^Y139^, A4^W160^/McpX^W161^, A4^Y176^/McpX^Y177^, and A4^D205^/McpX^D208^) occupy similar positions and orientations (Fig. 5c-d). However, residues R116, T145, N178, and N180, which are also important for uracil binding, are not conserved in McpX, indicating that these residues may provide specificity for uracil binding (Fig. 5a, c, d and Extended Data Fig. 8a). Indeed, replacing these four residues with the ones from McpX abolished uracil binding and enabled binding of epinephrine, but not other compounds from our HGMT ligand library (Fig. 5e and Extended Data Fig. 8b-e).

Furthermore, comparison of the membrane-proximal binding modules showed conservation of all four key residues for acetate binding between dCache_1UR and dCache_1AA (PctA), but not in the dCache_1AM and dCache_1PU domains (Fig. 5f). Consistent with that, bound acetate was reported in the published structure of the membrane-proximal module of PctA, although the physiological significance of this binding remains unclear^14^ (Fig. 5g–h). However, sequence analysis of dCache_1UR homologs indicates that the dCache_1UR and SCFA-binding motifs do not always co-exist (Supplementary Data 3).

### Short-chain carboxylic acids are ubiquitous chemotactic signals and valuable metabolites for gut bacteria

Given that the majority of ligands identified in our screen targeted bacterial chemoreceptors carrying Cache superfamily domains, we next focused on gut bacteria that contain chemoreceptors of this superfamily. According to MiST4.0 database^35^, 12 out of the 20 organisms from our initial set contain chemoreceptor genes, and 11 of these possess Cache-type chemoreceptors (Supplementary Data 1, 2). Overall, Cache-type chemoreceptors in these organisms account for over 30% of all transmembrane chemotaxis receptors (Extended Data Fig. 9), with the 4HB and MASE-type domains contributing the other two-thirds. Notably, MASE-type domains were not part of this study because they are characterized by multiple transmembrane regions and lack a clearly structured extracytoplasmic domain^36^.

To better understand the sensory repertoire of these receptors in the studied gut bacteria, we integrated experimentally identified ligand specificities with predictions based on sequence alignment (Fig. 6a and Extended Data Fig. 4). Altogether, we could assign ligands to 50% of the Cache family domains, including all sCache_2 domains, most of the sCache 3_3 domains, and approximately one-third of the dCache_1 domains (Extended data Fig. 9). Among the classes of metabolites, carboxylic acids were particularly prevalent chemotactic stimuli (Fig. 6b). Most common appear to be the sensors for L-lactate and formate, being carried by the majority of the studied organisms, strongly suggesting physiological importance of locating sources of these two metabolites for motile gut bacteria. A particularly striking example is *R. hominis* that is predicted to carry sensors for both L-lactate and pyruvate, as well as two sensors for formate. In contrast, fewer bacteria possess Cache sensors for the commonly studied and more abundant SCFAs (acetate, propionate, and butyrate). Importantly, the chemoreceptor *K*_D_ values for different intestinal metabolites fall within their known natural concentration ranges, and the receptor affinities thus appear to inversely correlate with the metabolite concentration in the gut, with highest receptor affinities for low-abundance metabolites and only low affinities for highly abundant SCFAs (Supplementary Table 3).

**Fig. 6.**
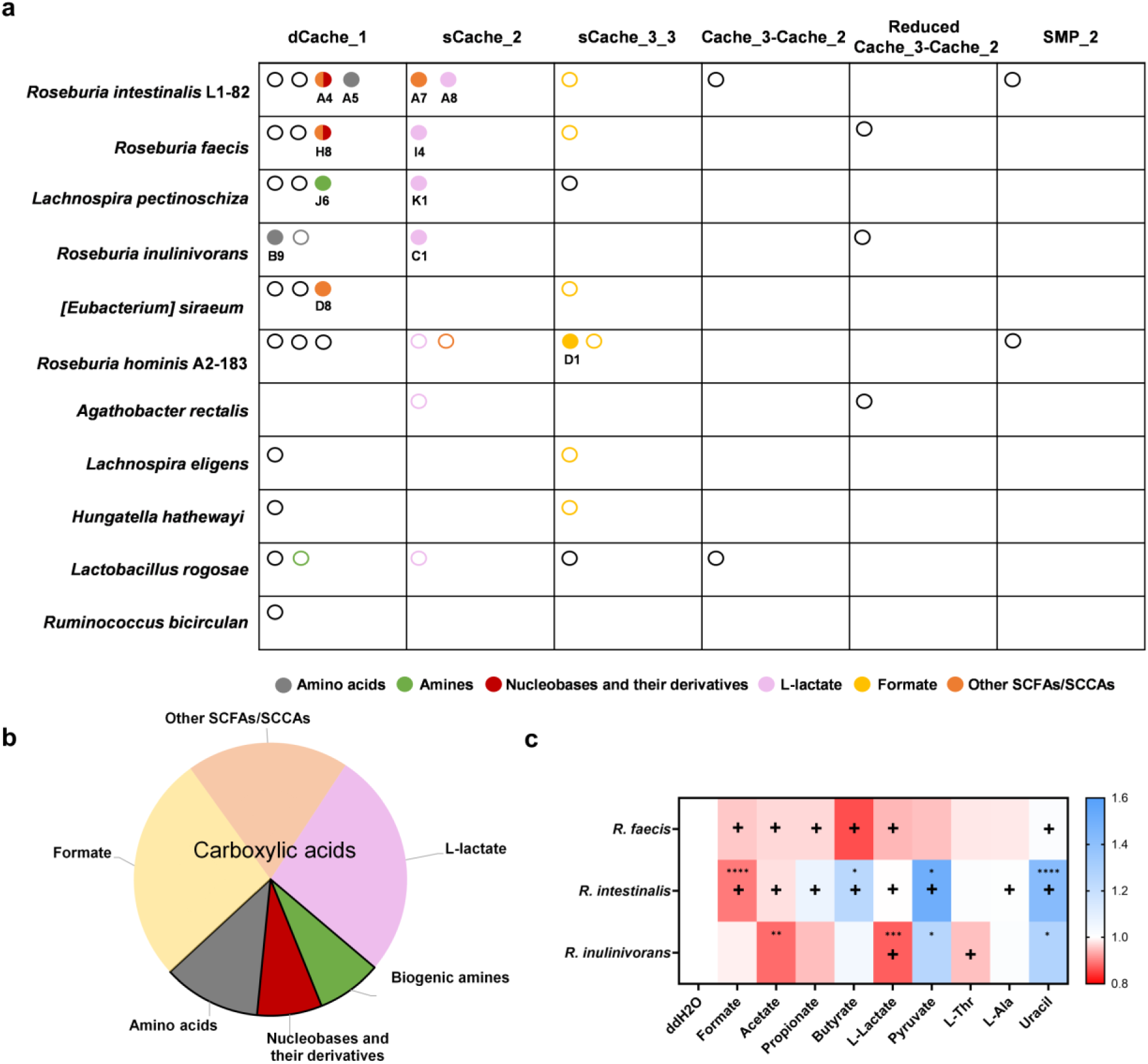
Overview of the chemotactic signals for the Cache superfamily domains of Clostridia in the gut microbiome. **a**, Chemotactic stimuli profiles of Cache domains in the indicated gut bacteria, identified using the experimental and bioinformatic analyses. Cache domains are categorized based on the domain family. Each LBD is represented by an open circle (predicted stimulus) or a filled circle (experimentally identified stimulus). Colors indicate distinct classes of ligands. **b**, Composition of identified and predicted chemoeffectors, categorized by classes of chemical compounds, as indicated. L-lactate, formate, and other short-chain fatty acids (SCFAs)/short-chain carboxylic acids (SCCAs) are collectively classified as carboxylic acids. **c**, Heat map showing the effects of addition of 20 mM of indicated compounds (10 mM for uracil) on growth of the selected *Roseburia* species in 50% YCFA medium (see Extended data Fig. 10 for respective growth curves), quantified as the time required to reach the maximal growth rate (t(μ_max)). The symbol “+” indicates the indicated strain possesses the identified or predicted chemoreceptor responsive to the corresponding compound. Data represent mean values from three biological replicates. *p* values (\**p* ≤ 0.05, \*\**p* ≤0.01, \*\*\**p* ≤0.001, and \*\*\*\**p* ≤0.0001) are assessed from a two-tailed unpaired *t*-test.

Such correlation could be consistent with the presumed primary physiological function of bacterial chemotaxis, namely in locating sources, and thereby enhancing acquisition, of limiting nutrients^17,18,37^. Hence, we directly evaluated the impacts of major identified chemoeffectors on the growth of three species, *R. intestinalis* L1-82, *R. faecis*, and *R. inulinivorans*. Although no straightforward correlation was observed between the growth effect of a specific metabolite and the presence of its chemoreceptor, most carboxylic acids, except pyruvate, promoted growth of at least one of the tested species (Fig. 6c and Extended data Fig. 10). The most significant impact on growth resulted from supplementation of the medium with formate or L-lactate, and it was observed for strains that possess respective receptors, supporting the hypothesis that chemotaxis toward carboxylic acids may be because of their nutritional value to the gut commensals. In contrast, the effect of supplementation with uracil was surprisingly growth-inhibitory, and the most significant impact was again observed for *R. intestinalis* L1-82 that possesses the corresponding chemoreceptor.

## Discussion

The gut microbiome is a complex community that relies on the extensive exchange of metabolites and signals among microorganisms and between microbes and the host^38,39^. However, it remains largely unknown which signals are recognized by diverse extracytoplasmic sensory domains that provide inputs for signal transduction pathways in gut bacteria. Previous systematic efforts on identifying ligands of bacterial sensors centered on individual types of sensory domains^31–33,40^ or on several individual model organisms, primarily pathogens^7,26,41–43^. Here we instead pursued a habitat-centric approach, assessing ligand specificities for a library of 116 LBDs from the recently established dataset of ∼17000 sensory domains of the human gut commensal bacteria^21^. We primarily focused on the bacterial class of Clostridia, since this class is highly prominent and important in the human gut microbiome^22^, and many of its members are motile and therefore possess all three major types of bacterial transmembrane sensors, including MCPs, HKs, and DGCs.

Collectively, our screening of these LBDs against a library of metabolites known to be present in the gut identified 34 receptor-ligand interactions, revealing distinct patterns in ligand distribution between the functional and structural families of bacterial sensors. Although the LBD library had nearly equal proportions of folded MCP and HK LBDs available for ligand binding assays, nearly all identified ligands were MCP-specific. Ligands could be mapped to 12 of 35 MCP LBDs but only to one of 33 HK LBDs. A possible explanation for this notable difference might be that most of the tested ligands are potential nutrients. Bacterial chemotaxis receptors are believed to preferentially recognize nutrients favored by particular bacteria, facilitating foraging and movement toward optimal growth environments^17,18,37,40^. Indeed, a recent compilation suggests that three-quarters of all known bacterial chemoattractants serve as growth substrates for the respective bacteria^44^. In contrast, two-component systems commonly regulate diverse stress responses^45^, and might therefore recognize signaling molecules other than nutrients. Consistent with this distinction between the ligand repertoire of these different functional classes of receptors, there are very few nutrients among the previously identified HK LBD ligands^40^. The only ligand identified for a sensory HK was indole, which is a ubiquitous bacterial signaling molecule rather than a nutrient^46,47^. Of note, despite a potentially widespread importance of indole as a bacterial signaling molecule across a range of natural environments, including the human GI tract^47,48^, HK B6 from *C. mitsuokai* is the first bacterial sensor to date with biochemically demonstrated indole binding to its extracytoplasmic sensory domain. Another sCache domain from *E. coli* BaeS HK has also been shown to respond to indole, although the underlying mechanism remains unknown^49^.

This observed difference in the LBD coverage between MCPs and HKs was even more surprising, as HKs contained a significantly larger proportion of Cache superfamily LBDs, to which all the characterized LBDs in our study belonged. In contrast, no ligands could be identified for the eleven folded 4HB domains that were included in our screen. One possible reason for that could be the insufficient stability of the ligand binding pocket of purified 4HB domains, given that they commonly bind ligands at the interface between the two monomers^50^, despite some exceptions^51^. This is different from LBDs from the Cache superfamily, which have one or two well-defined ligand-binding module(s) within the LBD monomer^52^. Nevertheless, both our control experiments with the 4HB domains of *E. coli* MCPs and previous studies demonstrate that the ligand binding of 4HB domains can be characterized using TSA^26,53^. Alternatively, such lack of identified ligands might indicate that 4HB domains more commonly interact with the ligand-loaded solute binding proteins (SBPs) than with the ligands themselves. Indeed, while all four *E. coli* 4HB domain MCPs are known to interact with SBPs, only two have been shown to directly bind ligands^7^. Such SBP-mediated signaling could also provide an alternative explanation for the lack of identified HK ligands, since at least some of the Cache domains of HKs have been shown to bind SBPs^54^.

The spectrum of nutrient metabolites that were identified as MCP ligands in our study shows high diversity, encompassing amino acids, amines, and pyrimidines. The most prevalent ligands, however, were carboxylic acids, where we identified three different binding modes: (1) sCache_2 domains that bind lactate or pyruvate; (2) a dCache_1 domain that binds dicarboxylates at its membrane-distal module; and (3) dCache_1 domains (named here dCache_1UR) that bind SCFAs at their membrane-proximal and uracil at their membrane-distal modules.

This latter example is particularly intriguing because, although dCache domains harbor two putative ligand-binding modules^55^, only a few examples of ligand binding to both sites within the same sensor have been described experimentally so far^56–58^ Our structural and phylogenetic analysis of the dCache_1UR domain complex with two ligands allows us to draw several conclusions about the evolution of this subset of dCache_1 domains. We demonstrated that the uracil-binding pocket is closely related to that of dCache_1AM (amine) sensors, and its specificity could be altered from uracil to amine through several amino acid substitutions. Remarkedly, this alteration yielded a sensor for epinephrine, a human catecholamine hormone. Although its physiological relevance remains to be investigated, epinephrine is known to play an important role in host-microbiota interactions as a neurotransmitter^59^, our findings demonstrate that dCache_1 domains can easily evolve specificity for epinephrine. In contrast to the uracil-binding distal pocket, the SCFA-binding proximal pocket shows similarity to the proximal pocket of the amino acid sensor PctA, which was indeed previously co-crystalized with acetate^14^. Our analysis suggests that specificities of the two pockets are not strictly coupled, but the detailed evolution of these receptor modules requires further investigation. Further emphasizing evolutionary adaptability of bacterial LBDs, we demonstrate the specificity of sCache_2 domains can be converted from L-lactate to another carboxylic acid by a single amino acid substitution.

Finally, our results provide general insights into the chemotactic preferences of commensal bacteria in a particular environment, the human gut. Overall, we could identify or predict ligand specificities for 50% of the Cache-superfamily sensors in the studied commensal bacteria. Whereas the better-studied sensors for amines and amino acids appear to be only sparsely represented, the less-studied sensors for carboxylic acids are highly prevalent in these bacteria. This preference is different from the current chemoattractant spectrum of the known bacterial species, where the most common attractants are amino acids and peptides, benzenoids, and purines^44^, which points to the nutritional value of carboxylic acids in the gut.

Interestingly, within this group, the typical ligands are not the commonly studied and highly abundant SCFAs but rather L-lactate and formate. Despite their low concentrations in the gut, L-lactate and formate likely play important roles in gut microbiota. L-lactate is produced in significant amounts by the host^60^, likely forming gradients toward the gut epithelium. Moreover, both L-and D-lactate, as well as formate can be produced by many gut bacteria as products of the anaerobic mixed-acid fermentation^60–62^, thus likely creating local sources of these metabolites. The observed detection of both lactate enantiomers, but with the preference for L-lactate, indicates that both host- and bacteria-derived lactate might be important metabolites for the gut Clostridia. Indeed, both lactate and formate can have important roles in bacterial cross-feeding under anaerobic conditions, serving as nutrients and/or electron donors for anaerobic respiration^63–65^. These conclusions align with our observation observed effects on the growth of selected commensal bacteria, where L-lactate and formate yield the most significant growth enhancement among the tested chemoeffector metabolites. This observation is further supported by LBD affinities, with the highest affinities observed for the less abundant metabolites. Finally, besides serving as a nutrient, L-lactate secreted by the host cells might further serve as a sensory cue, enabling bacteria to orient in the GI tract, as was previously proposed for other host-derived signals^66^. Such multiple roles of L-lactate – as a nutrient, signaling molecule, and a chemoattractant – have been implicated in the infection by the gastric pathogen *Helicobacter pylori*^52,67^.

Interestingly, although DGCs are not usually considered to be nutrient sensors^16^, nucleobase derivatives and (or) SCFAs were also identified as ligands for both tested sensory domains of the DGCs. In the latter case, the DGC LBD belonged to the same dCache_1UR domain family as chemoreceptor sensors, demonstrating its recent evolutionary exchange among gut bacteria. However, the extent to which DGCs in commensal bacteria can sense nutrients remains to be further investigated across a broader set of transmembrane DGC sensors.

## Materials and methods

### Bacterial strains, plasmids, and growth conditions

Bacterial strains, plasmids, and primers used in this study are listed in Supplementary Table 4. For molecular cloning and protein expression, *E. coli* strains were grown in Luria broth (LB; 1% tryptone, 0.5% yeast extract, and 1% NaCl) at 37 °C with shaking. For chemotaxis and FRET experiments, *E. coli* strains were grown in tryptone broth (TB; 1% tryptone and 0.5% NaCl) at 34 °C with shaking. When necessary, antibiotics were used at the following final concentrations: kanamycin, 50 µg/ml; ampicillin, 100 µg/ml; and chloramphenicol, 34 µg/ml. For the gut bacteria growth assays, detailed information is provided as a separate section.

### Construction of protein expression plasmids and chimeric chemoreceptors

The protein sequences of studied extracytoplasmic sensory domains (Supplementary Data 1) were back-translated into DNA sequences using EMBOSS Backtranseq^68^. Subsequently DNA sequences were codon-matched to *E. coli* using DNA Chisel^69^. During the process, the following enzyme recognition sites were excluded: BsaI, BsmBI, EcoRI, NotI, PstI, SpeI, and XbaI. Synthetic DNAs were ordered as fragments from Twist Bioscience. All protein expression plasmids were generated by Gibson Assembly. In brief, the periplasmic sensory domains with overlapping sequences of vector pET28a (+) were amplified by PCR using the synthetic DNA fragments as templates. The resulting fragments were assembled into the linearized vector pET28a (+) (digested by NdeI and BamHI) using in-house generated Gibson Assembly mix^70^. Single amino acid mutations were generated using the Q5 site-directed mutagenesis kit (New England BioLabs) following the manufacturer’s instructions. All constructs were verified by external Sanger DNA sequencing services.

To construct chimeric chemoreceptors, the amplified LBD fragments containing the overlapping sequences of vector pKG116 and random linkers were cloned into the linearized vector pKG116 digested by NdeI and BamHI. After cloning, the functional chimeras were selected from a library of the LBD [1-X]-XXXXX-Tar [203-553], which contains a five-amino acid random linker between the LBD and Tar signaling domain, as described previously^71^.

### Expression and purification of ligand binding domains

*E. coli* T7 Express strains (New England BioLabs) carrying the LBD expression plasmids were grown in LB medium supplemented with kanamycin at 37 °C until the optical density at 600 nm (OD_600_) reached 0.6. Isopropyl-β-D-thiogalactoside (IPTG) was added to induce protein expression at a final concentration of 0.1 mM. Growth was continued at 18 °C for 12 h and cells were collected by centrifugation. Proteins were purified by metal affinity chromatography using modified procedures for His GraviTrapTM column. Briefly, cell pellets were resuspended in binding buffer (20 mM sodium phosphate, 500 mM NaCl, and 20 mM imidazole, pH 7.4) supplemented with 0.2 µg/ml lysozyme, 1 mM MgCl_2_, 1 mM PMSF, stirred for 30 min at 4 °C and disrupted using an ultrasonic homogenizer followed by centrifugation at 20,000 x *g* at 4 °C for 30 min. Afterward, the supernatant was loaded into His GraviTrap^TM^ column previously equilibrated with binding buffer. Following two washing steps with binding buffer, proteins were eluted by elution buffer (20 mM sodium phosphate, 500 mM NaCl, and 500 mM imidazole, pH 7.4). Finally, the eluted protein fractions were dialyzed against dialysis buffer (10 mM sodium phosphate, 150 mM NaCl, 10% (v/v) glycerol, pH 7.0) and concentrated using Amicon Ultra-15 centrifugal filters. For proteins used in ITC experiments, an additional purification step was performed by size-exclusion chromatography (SEC) on an S200 XK16 column (Cytiva) using a buffer with the same composition as the dialysis buffer.

To remove the bound acetate and uracil from the A4 LBD, the protein was purified under previously established conditions with the addition of two extra washing steps during the affinity purification. Briefly, the cleared lysate was loaded into 1 ml HisTrap HP column (GE Healthcare), followed by three sequential washing steps prior to elution: (1) 20 column volumes (CV) of binding buffer, (2) 10 CV of washing buffer, and (3) 20 CV of binding buffer. To evaluate the best conditions for removing the pre-bound ligands, different washing buffers were tested: (1) 20 mM sodium phosphate, 2 M NaCl, pH 7.4; (2) 20 mM sodium phosphate, 3 M Urea, pH 7.4; (3) 20 mM sodium phosphate, 5 M Urea, pH 7.4; and (4) 20 mM sodium phosphate, 7 M Urea, pH 7.4. The purified protein was analyzed using HPLC to determine the amount of uracil present in the sample. For subsequent experiments, the washing solution containing 5 M urea was selected.

### HPLC analysis

HPLC analysis was conducted using an Agilent 1,100 Series System (Agilent Technologies) equipped with a Metrosep A Supp5 – 150/4.0 column (Metrohm). 10 µl of each sample were injected. Nucleotides were eluted at a flow rate of 0.6 ml/min with 90 mM (NH_4_)_2_CO_3_ at pH 9.25 and detected at the wavelength of 260 nm. The sample was prepared as follows: 150 μl of chloroform was added to a 50 μl solution containing 10 μM of purified protein. The mixture was then vigorously agitated for 5 s, heated at 95 °C for 15 s, and immediately snap-frozen in liquid nitrogen. The thaw samples were centrifuged (13,000 r.p.m. for 20 min at 10 °C), and the aqueous phase was transferred to an HPLC vial for analysis. A 50 μl solution containing 1 mM uracil was used as standard and treated using the same procedure as described above. The resulting chromatograms were plotted using GraphPad Prism v8.4.3, and the uracil standard data was transformed using Y=Y/100 to facilitate the visual comparison with sample results.

### Thermal shift assays

Thermal shift assays were performed in 384 microtiter plates using a Bio-Rad CFX384 Touch^TM^ Real-Time PCR instrument. The tested compounds are derived from two HGMT (human gut metabolites) plates that are preconfigured 96 well plates containing about 150 different chemical compounds, as demonstrated in Supplementary Table 2. All compounds were dissolved in H_2_O at the final concentration of 20 mM and pH 7.0. Each 25 μl assay mixture consisted of 20.5 μl of purified protein (30 -100 μM), 2 μl of SYPRO™ Orange (Invitrogen) at a 5x concentration, and 2.5 μl of chemical solution from HGMT plates, resulting in 2 mM final concentration of tested ligands, unless otherwise indicated. Samples were gradually heated from 23 °C to 95 °C at a rate of 1 °C per min. The unfolding of proteins was tracked by detecting changes in fluorescence. This process enabled the determination of the midpoint of the protein unfolding transition, also known as the melting temperature (Tm), by utilizing the first derivative values of the raw fluorescence data. Data analysis was conducted using Bio-Rad CFX Manager 3.1 software.

### ITC measurements

Ligands and proteins were diluted in a buffer containing 10 mM sodium phosphate, 150 mM NaCl, 10% (v/v) glycerol, pH 7.0. The purified proteins were titrated in the sample cell at a final concentration of 25-91 µM each (Supplementary Table 5). The protein concentrations were predetermined by absorbance at 280 nm. The ligands were placed in the titration syringe at a nominal concentration of 0.1 to 10 mM to saturate the protein sample during the titrations. All the measurements were performed at 25 °C with the instrument MicroCal PEAQ-ITC (©Malvern Panalytical) with a method consisting of 19 injections (first 0.4 µl, and the rest 2 µl each) and 150s of spacing. The raw data were processed with the MicroCal PEAQ-ITC Analysis Software using the “one binding site” model and plotted using GraphPad Prism v8.4.3.

### Competitive ITC measurements

Competitive ITC measurements were performed to investigate the interplay between ligand-binding events mediated by two distinct ligand-binding modules. The detailed protocol was modified from the previous publication^72^. Briefly, 66-69 µM of Apo-A4 LBD protein was incubated in the buffer containing 10 mM sodium phosphate, 150 mM NaCl, 10% (v/v) glycerol, pH 7.0, and supplemented with either 700 µM uracil or 3 mM acetate. For titration, the same buffer as for protein preparation was used, containing either 700 µM uracil or 3 mM acetate. The same conditions were used for the measurement and data analysis, as described above.

### Soft-agar chemotaxis assays

The chemotaxis assay was conducted on semi-solid minimal A agar plates (0.25% (w/v) agar, 10 mM KH_2_PO_4_/K_2_HPO_4_, 8 mM (NH4)_2_SO_4_, 2 mM citrate, 1 mM MgSO_4_, 0.1 mg/ml of thiamine-HCl, 1 mM glycerol, and 40 μg/ml of a mixture of threonine, methionine, leucine, and histidine) supplemented with appropriate antibiotics and inducers. After solidification, 200 µl aliquots of 100 mM chemical solutions were applied as a line to the center of the plate and incubated at 4 °C for 16 h to form a chemical gradient. The chemoreceptor-less *E. coli* cells expressing the chimera as a sole chemoreceptor were plated approximately 2.5 cm away from the line where the chemical was applied, and plates were incubated at 30 °C for 24 - 48 h.

To select a functional chimeric chemoreceptor with a random linker, the library was applied to a soft agar plate with D-glucose gradients for three rounds of selection. D-glucose is a nonspecific chemoattractant that can be sensed via the phosphotransferase system (PTS) which transmits signals to the cytoplasmic region of chemoreceptors independently of the sensory domain^26^. Strains that migrated furthest in the D-glucose gradient were re-inoculated on a new plate for the next round of selection. After three rounds of selection, chimera expression plasmids for the best-chemotactic cells were isolated and their linkers were identified by external Sanger DNA sequencing services.

### FRET measurements

FRET measurements were performed as described previously^25,73^. Chemoreceptor-less *E. coli* strains VS181 with the plasmids encoding chimeric chemoreceptor and CheY-YFP/CheZ-CFP FRET pair were grown in 10 ml TB medium supplemented with appropriate antibiotics and inducers (50 µM IPTG and 1-2 µM sodium salicylate) at 34°C and 275 r.p.m. Cells were then harvested at OD_600_ of 0.5 by centrifugation and washed twice with tethering buffer (10 mM KH_2_PO_4_/K_2_HPO_4_, 0.1 mM EDTA, 1 μM methionine, 10 mM sodium lactate, pH 7.0), whereas the responses to lactate were measured using tethering buffer without sodium lactate. For microscopy, the cells were attached to the poly-lysine-coated coverslips for about 10 - 15 min and mounted into a flow chamber that was maintained under a constant flow of 0.3 ml/min of tethering buffer using a syringe pump (Harvard Apparatus) that was also used to add or remove compounds of interest. Given pH is a prevalent chemotactic stimulus, the pH value of all tested compounds was adjusted to 7.0. FRET measurements were performed on an upright fluorescence microscope (Zeiss AxioImager.Z1) equipped with photon counters (Hamamatsu). Finally, the fluorescence signals were recorded and analyzed as described previously^73^. D-glucose was routinely used as a nonspecific chemoattractant to assess the activity of hybrid chemoreceptors.

### Protein structure manipulations, computational docking, and protein sequence alignments

The structures of the target proteins (M8, H8, D8, and K1) were built using AlphaFold3^74^. The crystal structures of the pyruvate sensor (PDB ID 4EXO), PctA (PDB ID 5T65), and McpX (PDB ID 6D8V) were taken from the RCSB protein data bank^27^. Comparative analysis of solved and modeled protein structures was performed using PyMOL 3.0 and/or ChimeraX 1.8^75^. Figures were generated by ChimeraX 1.8. For the *in-silico* docking, ligands were downloaded from PubChem database^76^ in SDF format. DiffDock^77^ was used to computationally dock ligands to Cache domain models using the default settings. Docking results were viewed and interpreted using ChimeraX 1.8. Protein sequence alignments were built using the L-INS-I algorithm of the MAFFT package^78^ with default setting and visualized in Jalview^79^ to identify the presence of corresponding ligand-binding motif, employing the previously well-characterized motif containing sensors as a target.

To identify dCache_1UR homologs, two iterations of PSI-BLAST searches were initiated against the RefSeq database (release 226) with a maximum number of target sequences of 20,000 using the dCache_1 domain of the uracil sensor A4. Protein sequence regions corresponding to the dCache_1 domain were extracted, with sequence redundancy reduced at 100% identity, and a multiple sequence alignment was constructed using the MAFFT (v. 7.490)^78^ L-INS-I algorithm. Uracil and the putative SCFA motif residues were then identified and tracked on the alignment. Motif variant identification and taxonomy information retrieval were performed using a custom Python script.

### Gut bacteria growth measurements

Gut bacteria were cultivated at 37 °C under anaerobic conditions in a vinyl anaerobic chamber (COY) inflated with a gas mix of approximately 10% carbon dioxide, 88% nitrogen and 2% hydrogen. Prior to the measurement, frozen glycerol stocks were streaked for single colonies onto YCFA agar plates prepared according to the YCFA medium (ID1611) recipe provided by MediaDive^80^, and incubated at 37 °C. A single colony was inoculated in 1 ml YCFA medium and grown at 37 °C with constant shaking.

To monitor the growth effect of chemical compounds, bacteria growth assays were carried out using 50% YCFA medium, which was prepared by diluting the original YCFA medium with an equal volume of distilled water (ddH_2_O). All test compounds were prepared at a concentration of 20 mM, except for uracil, which was tested at 10 mM due to its poor solubility. Pre-cultures were diluted 1:100 into 500 µl of 50% YCFA medium supplemented with the respective test compound in a 48-well plate. Afterwards, growth was measured as OD_600_ in a plate reader (Infinite M200 Pro, Tecan) at 37 °C with constant shaking at 200 r.p.m. To correct for the medium turbidity, raw growth curves were first normalized with inoculum OD (blank OD). Growth plots were then generated using the GraphPad Prism (v. 10.1.0) and quantification of growth parameters was done using QurvE^81^ (non-parametric model). Statistical analyses were performed with GraphPad Prism (v. 10.1.0).

### Crystallization and structure determination of the A4 domain

Crystallization was performed using the sitting drop vapor diffusion method at 20 °C in 0.5-0.75 µl drops. The drops consisted of protein and precipitant solutions mixed at 1:1 or 1:2 ratios. The protein solution was prepared by incubating 35 µM of freshly purified His6-A4 with 700 µM uracil and 3 mM propionate for 20 min on ice. Then the protein solution was concentrated to 850 µM using 10 kDa cut-off Amicon Ultra Centrifugal Filter tubes (Millipore). A total of 384 crystallization conditions were screened using the JCSG Core Suite I-IV (Qiagen). Crystals were obtained in a solution containing 0.2 M potassium fluoride supplemented with 20% (w/v) PEG3350. Prior to data collection, crystals were flash-frozen in liquid nitrogen using a cryo-solution consisting of mother liquor supplemented with 20% (v/v) glycerol. Diffraction data were collected under cryogenic conditions at the European Synchrotron Radiation Facility (Grenoble, France) at beamline ID23-2. Data were processed with XDS and scaled with XSCALE^82^. The structure was determined by molecular replacement with PHASER^83^, using the AlphaFold3 model of A4 LBD. Manual model building was performed in COOT^84^, followed by the refinement with PHENIX^85^. Figures were prepared using ChimeraX 1.8. Crystallization data collection and refinement statistics are given in Supplementary Table 6. Structure coordinates and structure factors of A4 LBD have been deposited in the Protein Data Bank under the accession code (PDB ID): 9HVJ. Topology analysis of the A4 LBD was performed using tools from PDBsum^86^ and ChimeraX 1.8.

### Phylogenetic tree analysis

The multiple sequence alignment of dCache_1 protein sequences prepared to build the tree was edited using an alignment trimming tool, trimAl (v. 1.4.1)^87^: positions in the alignment with gaps in 10% or more of the sequences were removed unless this leaves less than 60%. In such case, the 60% best (with fewer gaps) positions were preserved. The amino acid replacement model for the set of protein sequences was determined running ProtTest (v. 3.4.2)^88^. The best model was found to be LG with gamma distribution of rate variation across sites in combination with the empirical state frequencies (LG + G + F). Using the determined amino acid replacement model, a phylogenetic tree was constructed using a Bayesian inference algorithm implemented in MrBayes (v. 3.2.7a)^89^. Metropolis-coupled Markov chain Monte Carlo simulation implemented in MrBayes was run with 3 heated and 1 cold chain, discarding the first 25% of samples from the cold chain at the “burn-in” phase. A total of 2,500,000 generations were run till the sufficient convergence was achieved (the average standard deviation of split frequencies is equal to or less than 0.01; in this case it was 0.008) with chain sampling every 1000 generations.

### Data availability

All data needed to evaluate the conclusions in the paper are present in the paper and/or the Supplementary Materials. The structure factors and coordinates have been deposited in the Protein Data Bank (PDB, www.rcsb.org) with the accession codes: 9HVJ.

## Acknowledgements

This work was partially supported by Max Planck Society (to V.S), Hessian Ministry of Higher Education, Research and the Arts (HMWK) –LOEWE research cluster “Diffusible Signals,” subproject A1 (to V.S.), National Institutes of Health Grant R35GM131760 (to I.B.Z.), Deutsche Forschungsgemeinschaft (DFG; Projektnummer 464366151) in the framework of the priority program “SPP 2330 – New Concepts in Prokaryotic Virus-host Interactions” (to G.B.).

## Author contributions

Conceptualization: W.X., V.S., and I.B.Z.

Investigation: W.X., E.J.K., V.M.G., T.S.K., D.S., and P.A.R.

Visualization: W.X., E.J.K., and V.M.G.

Supervision: V.S., I.B.Z. and G.B.

Writing—original draft: W.X. and V.S.

Writing—review & editing: W.X., V.S., E.J.K., V.M.G., I.B.Z., and G.B.

## Competing interests

The authors declare that they have no competing interests.

## Extended data

**Extended Data Fig. 1.**
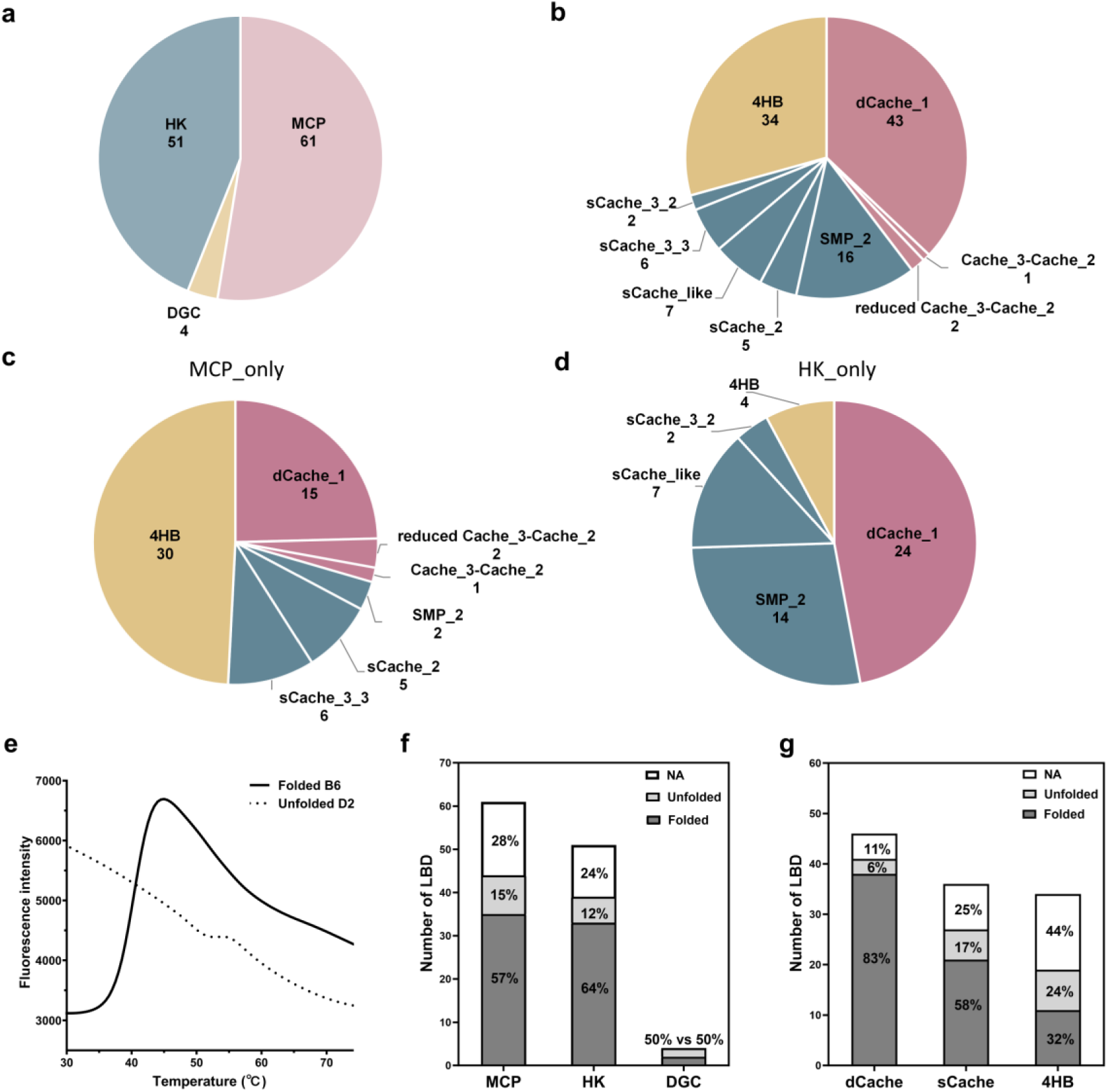
Functional and structural groups of the synthesized LBDs. **a**, **b**, Composition of the entire LBD library presented by the receptor type (a) and by the domain family (b), respectively. The numbers below each domain or receptor name indicate the number of LBDs in the respective subgroup. **c**, **d**, The proportion of LBDs from different domain families among all chemoreceptors (c) and histidine kinases (d). The underlying data are available in Supplementary Data 1. **e**, Representative melting curves of a folded (B6 LBD) and an unfolded domain (D2 LBD). **f**, **g**, Comparative analysis of protein thermal stability by receptor type (f) and domain family (g). Proteins that could not be expressed or purified, and thus could not be analyzed (NA) by thermal shift assays, are indicated in white; and purified proteins are categorized by thermal stability into unfolded and folded groups, shown in light gray and gray, respectively.

**Extended Data Fig. 2.**
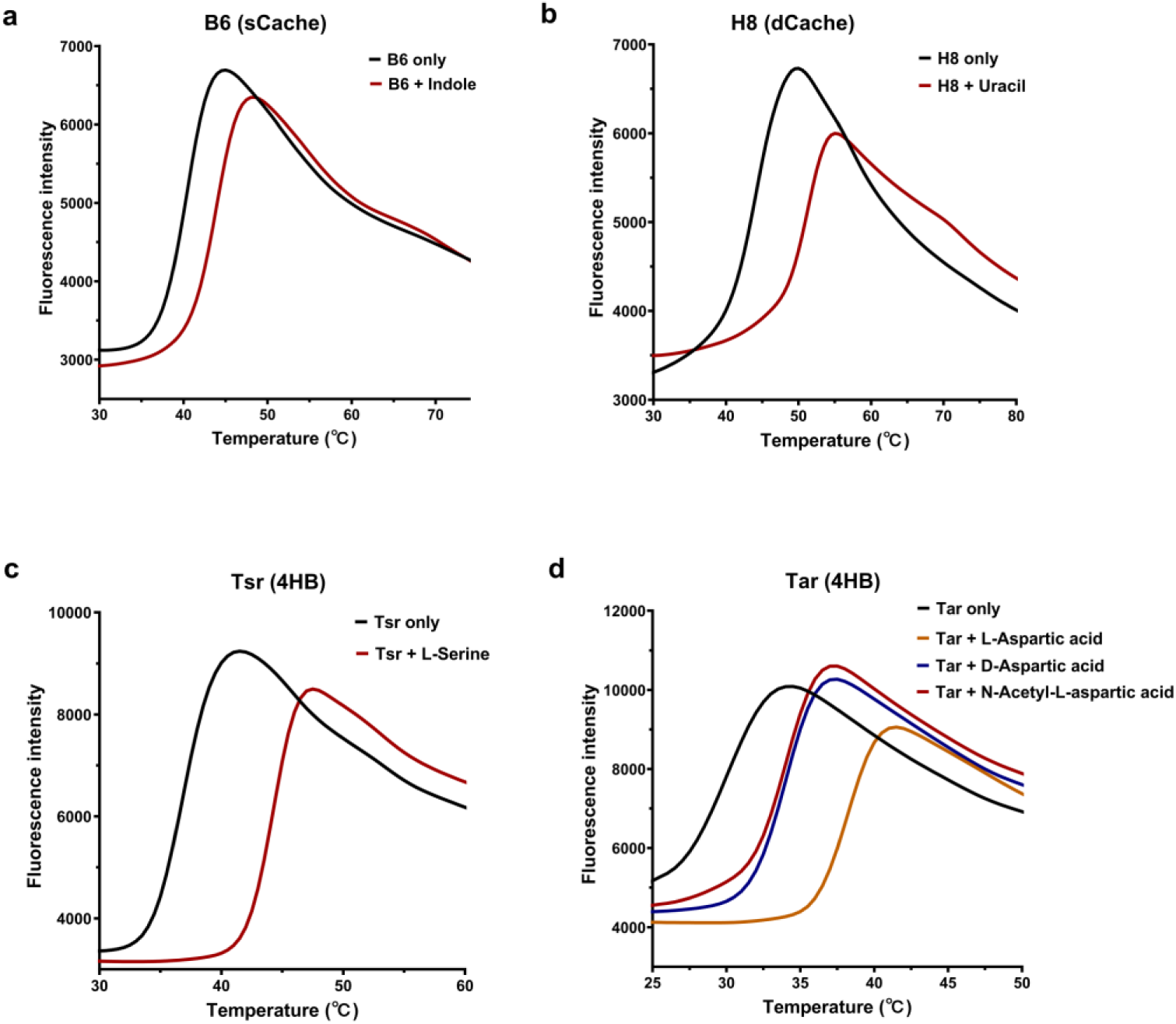
Melting curves of selected representative sensory domains. **a**, **b**, Representative thermal shifts of sCache and dCache domains upon addition of 2 mM final concentration of indicated ligands. Thermal unfolding curves of the sCache domain B6 (a) and the dCache domain H8 (b) in the absence and presence of indole and uracil, respectively. **c**, Thermal shift assays of the 4HB-type LBD from *E. coli* Tsr chemoreceptor using two HGMT plates. The Tsr LBD specifically binds L-serine. **d**, Thermal shift assays of the 4HB-type LBD from *E. coli* Tar chemoreceptor using two HGMT plates. The Tar LBD exhibited a positive thermal shift upon stimulation with L-aspartic acid (orange), D-aspartic acid (blue), and N-acetyl-L-aspartic acid (red).

**Extended Data Fig. 3.**
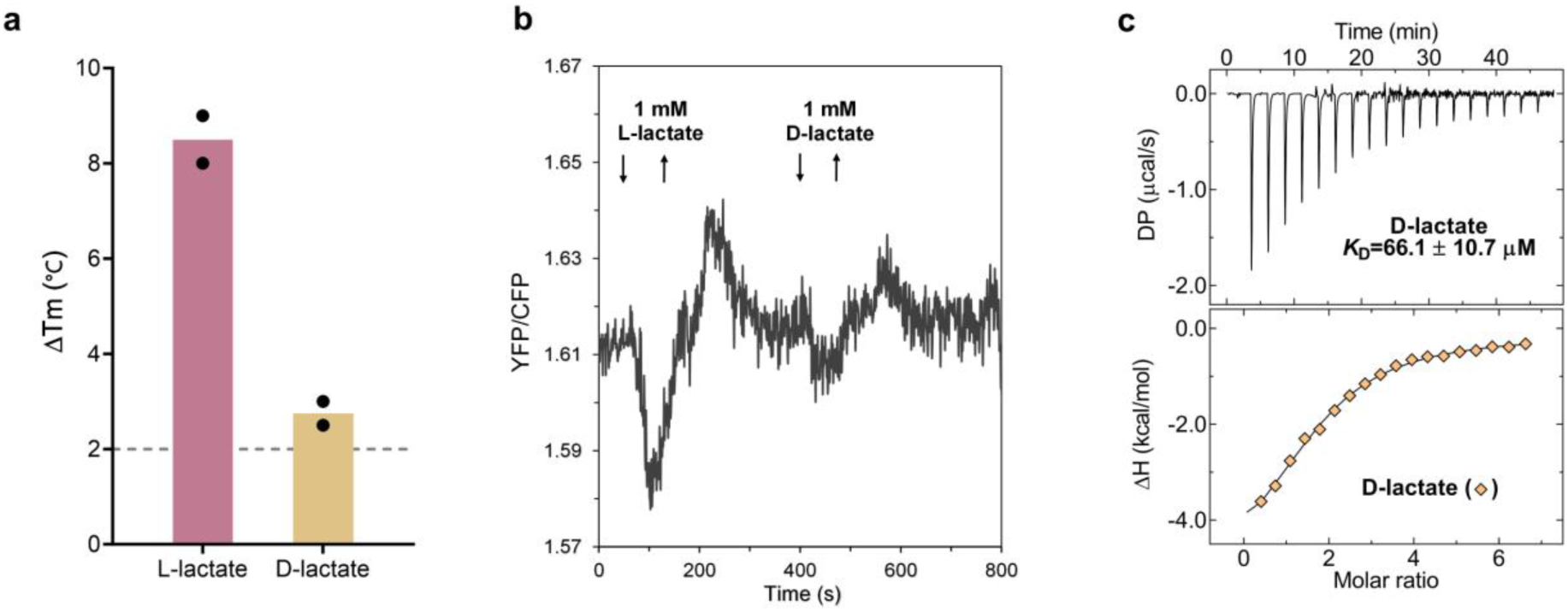
Characterization of K1 sensory domain binding and signaling response to D-lactate. **a**, Thermal shifts observed for the K1 sensory domain upon exposure to 2 mM of either L-lactate or D-lactate. Data represent the mean of two biological replicates (indicated by dots). **b**, FRET measurements of the K1-Tar chimera response to 1 mM L-lactate or D-lactate. Buffer-adapted *E. coli* cells expressing the CheZ-CFP/CheY-YFP FRET pair and the K1-Tar chimera as the sole receptor responded to stepwise addition (down arrow) and subsequent removal (up arrow) of stimui. **c**, Measurement of the binding affinity of K1 sensory domain to D-lactate using ITC. The upper panel shows raw titration data, and the lower shows integrated corrected peak areas of the titration data fitted using the “One binding site” model. The derived dissociation constant is shown in the upper panel. Further experimental details are provided in Supplementary Table 5.

**Extended Data Fig. 4.**
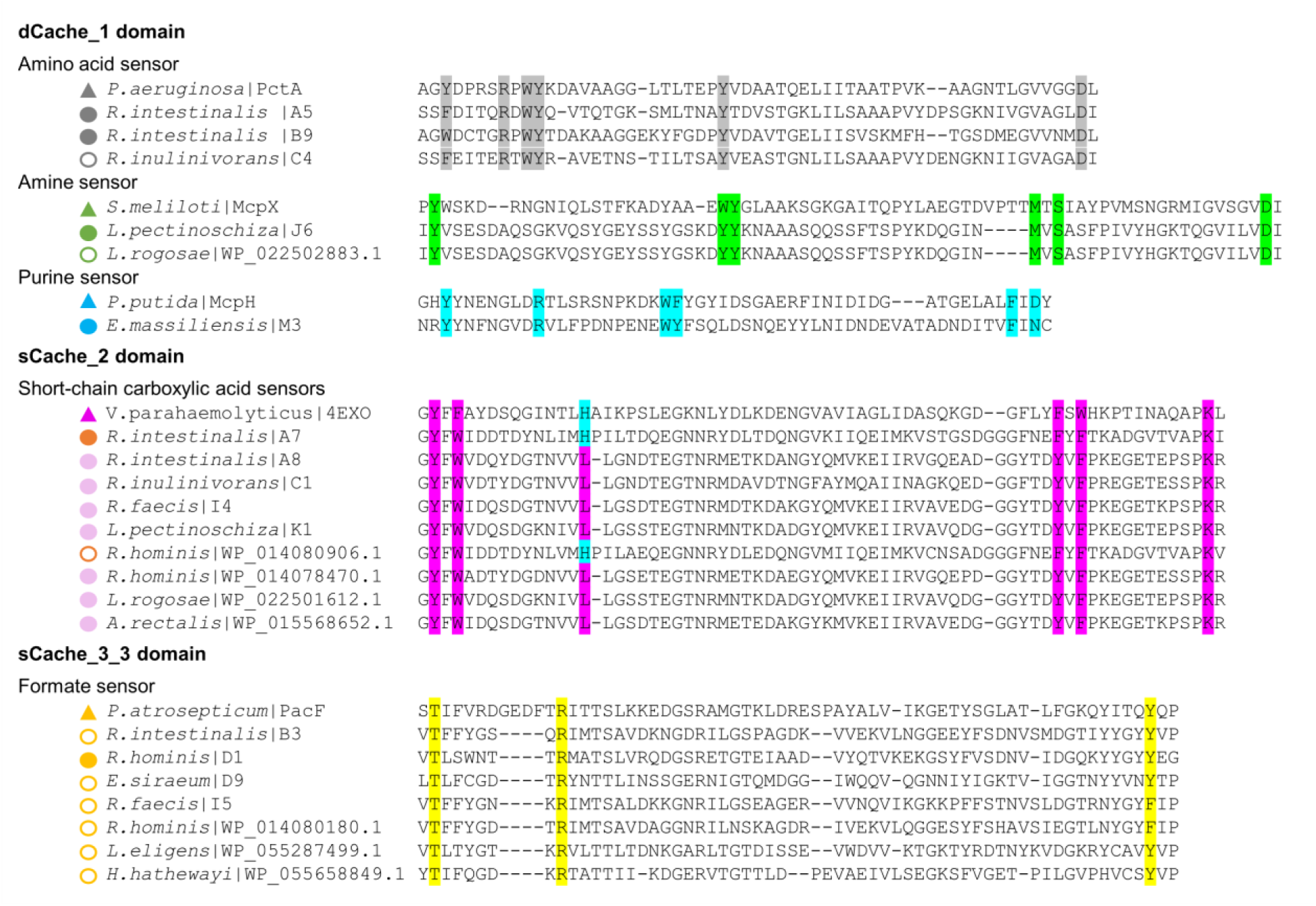
Multiple sequence alignments for LBDs containing characterized ligand-binding motifs. The key ligand-binding residues (motifs) were identified through sequence alignments with previously well-characterized sensory domains, including PctA (an amino acid sensor from *Pseudomonas aeruginosa*), McpX (an amine sensor from *Sinorhizobium meliloti*), McpH (a purine sensor from *Pseudomonas putida*), 4EXO (a short-chain carboxylic acid sensor from *Vibrio parahaemolyticus*), and PacF (a formate sensor from *Pectobacterium atrosepticum*). These previously characterized LBDs are labeled with triangles. LBDs labeled with solid circles denote the newly experimentally identified sensors in this study, while those labeled with open circles denote putative sensors predicted to bind the indicated ligands. Distinct sensor clusters are color-coded based on ligand specificity: gray for amino acid, green for amines, blue for purines, orange for pyruvate, pink for lactate, and yellow for formate.

**Extended Data Fig. 5.**
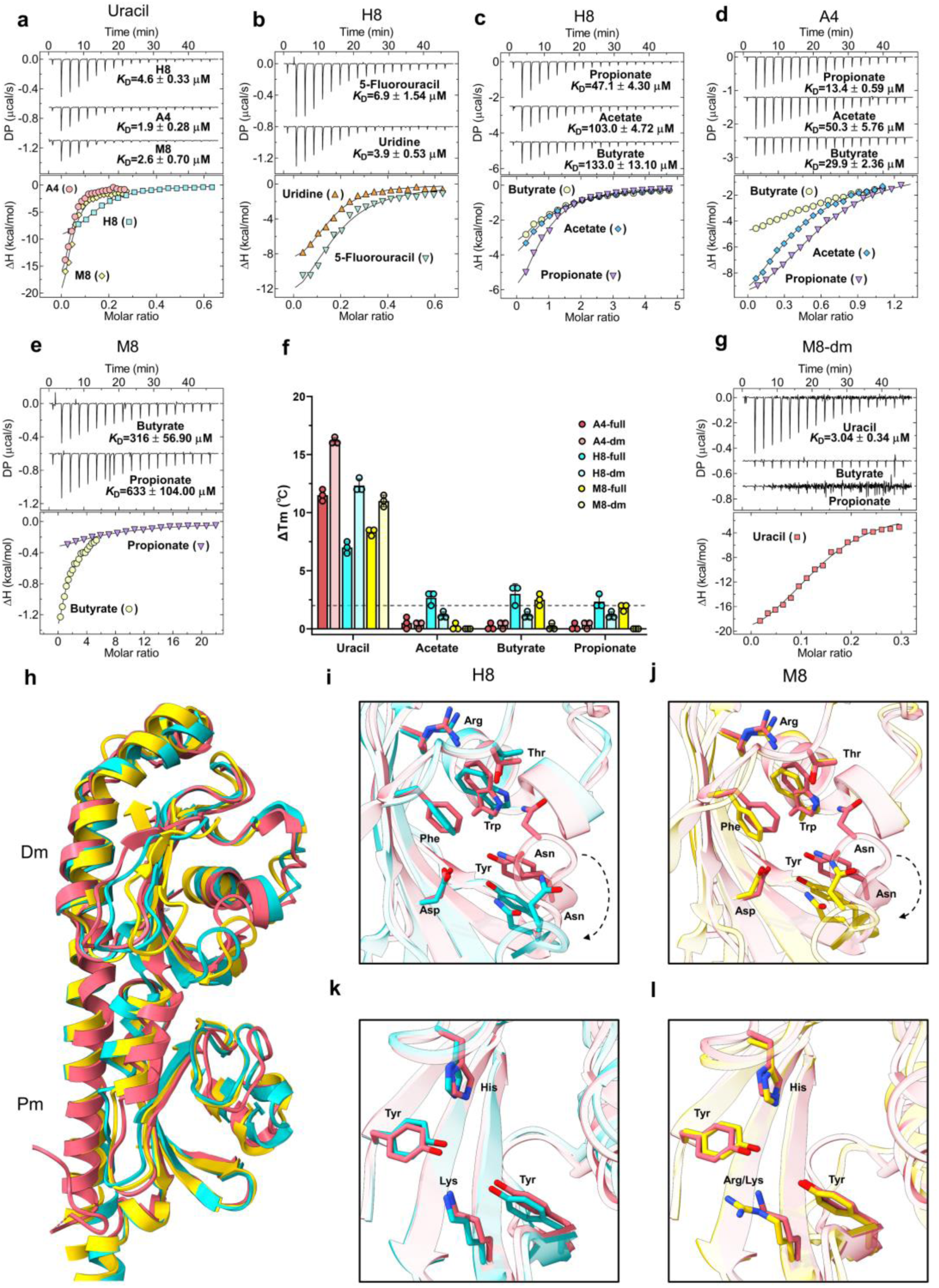
Characterization of uracil and short-chain fatty acid binding to dCache_1UR domains. **a**, ITC studies of uracil binding to the H8 LBD, M8 LBD, and A4 LBD. Upper panels: Raw titration data. Lower panels: Integrated, dilution heat corrected, and concentration normalized raw data. The lines are the best fits using the “One binding site” model, with the derived dissociation constants (*K*_D_) being indicated. **b**, ITC studies of H8 LBD with uridine and 5-fluorouracil. **c-e**, ITC studies of H8 LBD (c), A4 LBD (d), and M8 LBD (e) with the indicated short-chain fatty acids (SCFAs). **f**, Thermal shift measurements for the full-length (full) and membrane-distal module (dm) of H8, M8, and A4 in the presence of 2 mM concentrations of the indicated compounds. Data are presented as the mean ± standard deviation from three replicates, with each point indicates an individual measurement. **g**, ITC studies of M8-dm with uracil, butyrate, and propionate. Further experimental details are provided in Supplementary Table 5. **h**, The overall structural superimposition of the A4 structure with AlphaFold3 models of H8 and M8. Three uracil sensors are depicted in different colors: A4 - red, H8 - cyan, and M4 - yellow. **i**, **j**, Structural alignment of uracil binding pocket of A4 with the membrane-distal modules of H8 AlphaFold3 model (i) and M8 AlphaFold3 model (j). The amino acids in H8 and M8 corresponding to the uracil-binding key residues of A4 are labeled. **k**, **l**, Structural alignment of acetate binding pocket of A4 with the membrane-proximal modules of H8 AlphaFold3 model (k) and M8 AlphaFold3 model (l). The amino acids in H8 and M8 corresponding to the SCFA binding key residues of A4 are labeled. The dashed arrow indicates a loop displacement between the crystallographic structure of A4 and the AlphaFold3 models of H8 and M8.

**Extended Data Fig. 6.**
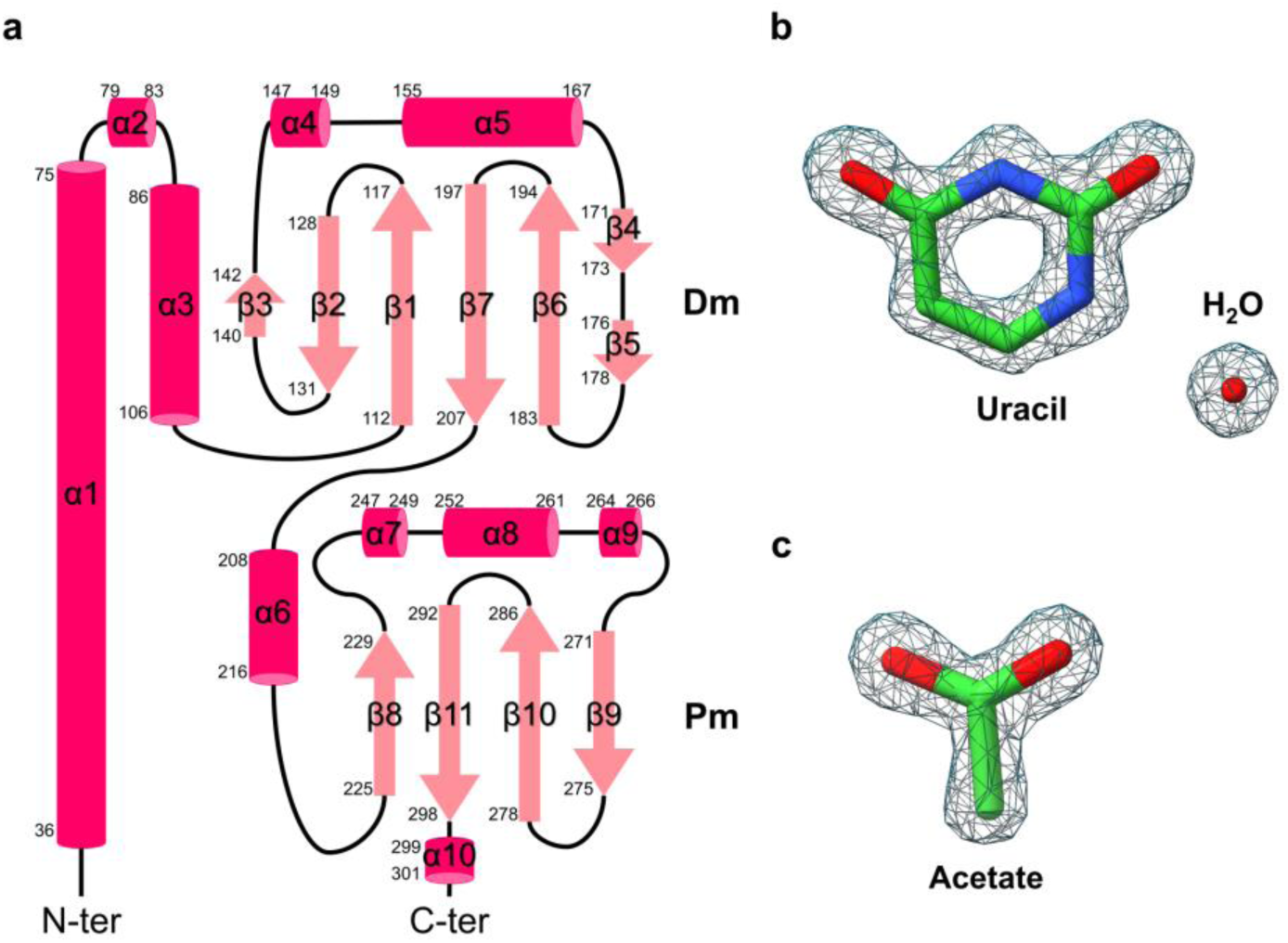
Topology of the A4 LBD and electron density of ligands. **a**, Topology of the A4 LBD secondary elements. The α-helices are represented by cylinders and the β-sheets by arrows. Dm and Pm stand for membrane-distal module and membrane-proximal module, respectively. **b**, **c**, 2mFo-DFc electron density maps of uracil and a water molecule (b), and acetate (c). 2mFo-DFc map (gray mesh) is contoured at 1.24σ.

**Extended Data Fig. 7.**
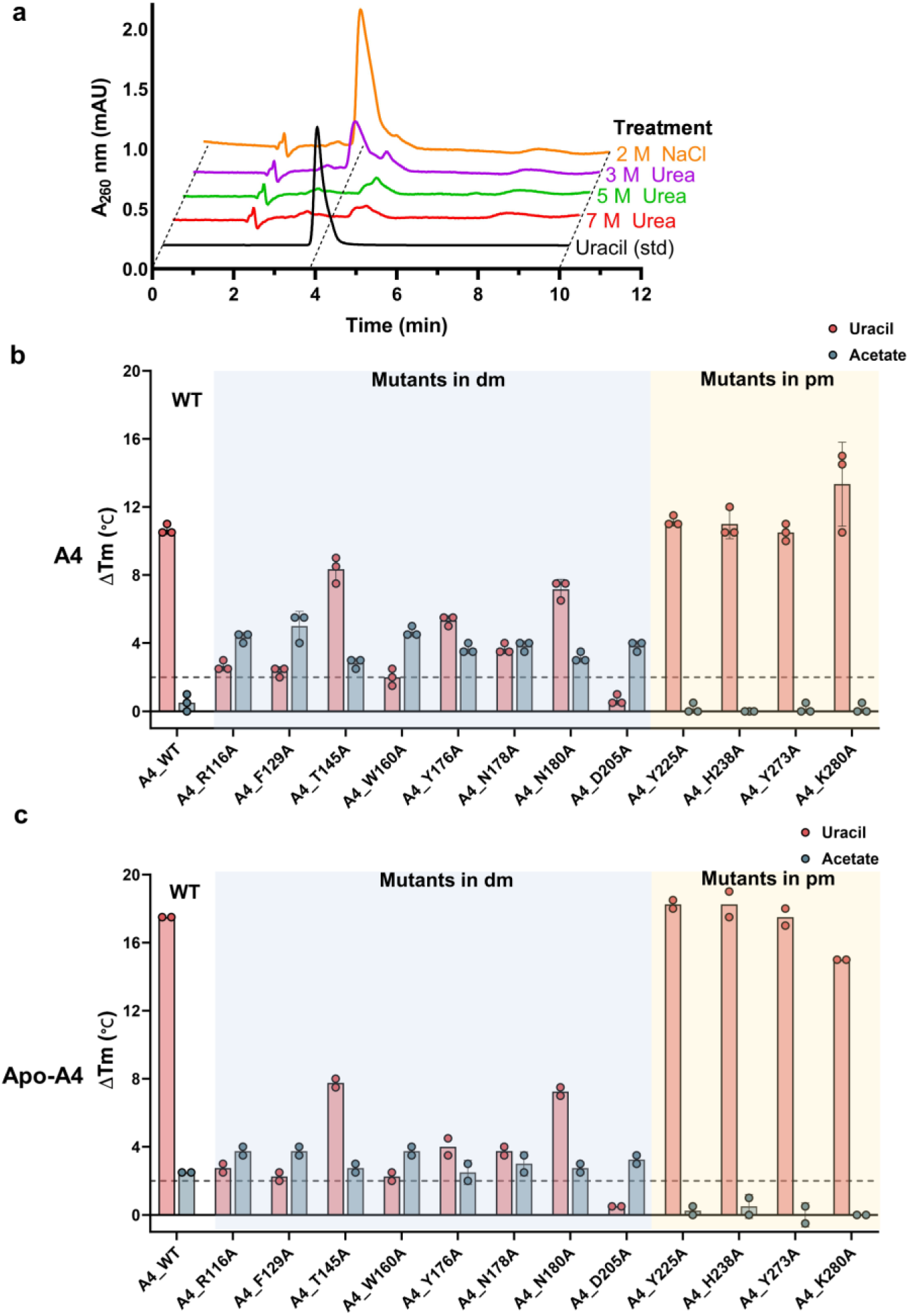
Experimental evaluation of the contributions of individual key amino acids in the A4 LBD to ligand binding. **a**, HPLC chromatogram of the A4 LBD after different treatments to remove endogenously bound ligands. A solution containing only uracil (black trace) was used as a standard. **b**, Thermal shift measurements for the indicated A4 LBD mutants in the presence of 2 mM uracil or acetate. The data represent the mean ± standard deviations of three biological replicates. **c**, Thermal shift measurements for the apo-A4 BD mutants in the presence of of 2 mM uracil or acetate. The data represent the means of two biological replicates. Data are categorized into three groups: wild-type A4 (WT), A4 with mutations in the membrane-distal module (mutants in dm; blue), and A4 with mutations in the membrane-proximal module (mutants in pm; yellow). Each data point represents an independent biological measurement. The gray dashed line indicates the threshold of 2°C used as a significance cutoff for ligand identification.

**Extended Data Fig. 8.**
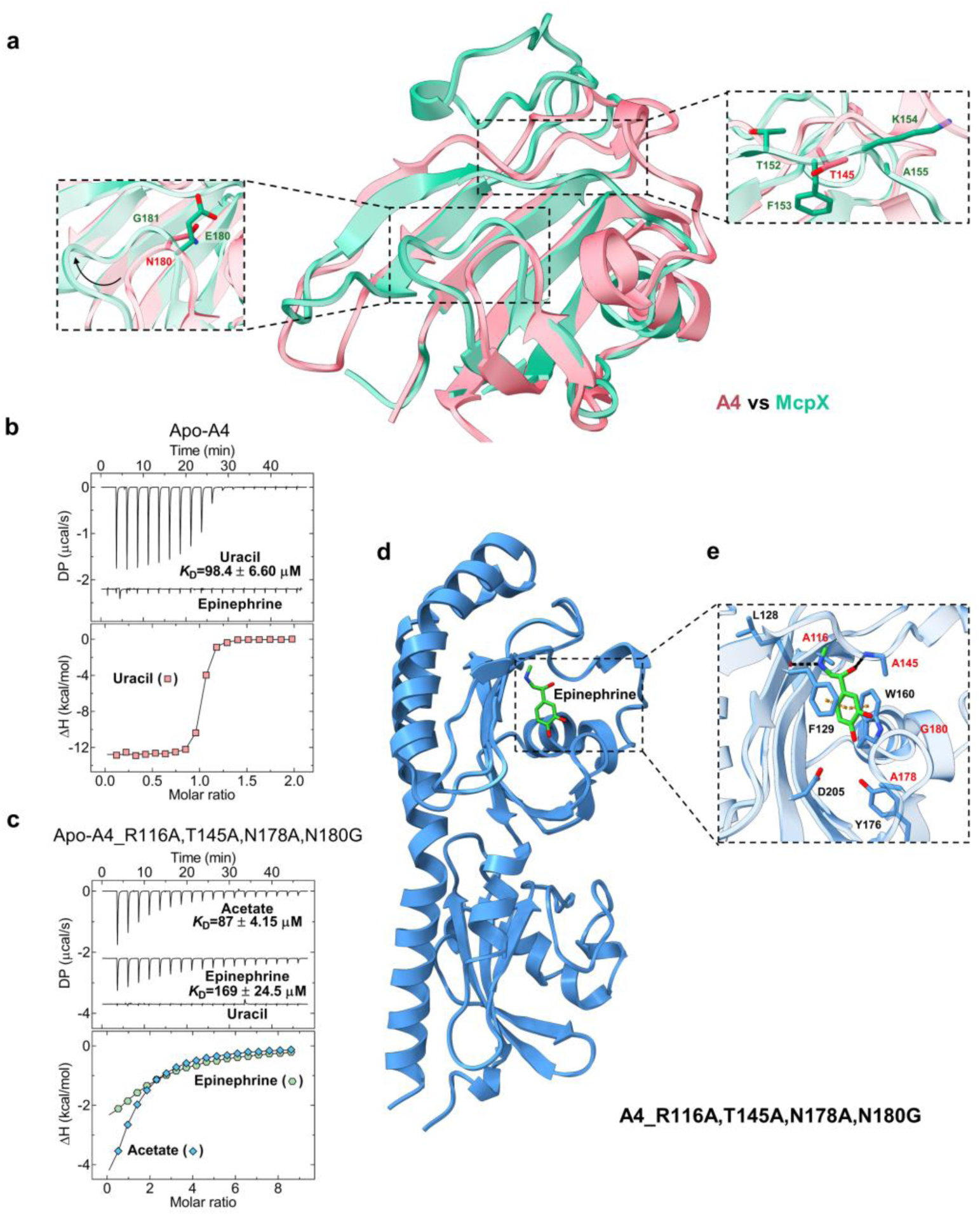
Binding of epinephrine to the A4_R116A, T145A, N178A, N180G LBD mutant. **a**, Structural superimposition of the membrane-distal module of the A4 (red) and McpX (green) LBDs. Close-up views highlight T145 and N180 in A4, along with their corresponding residues in McpX, as determined through structural (F153 and E180) and sequence alignments (A155 and G181). **b**, **c**, ITC measurements of Apo-A4 LBD (b) and Apo-A4_R116A, T145A, N178A, N180G mutant LBD (c) binding to uracil and epinephrine. The upper panel shows raw titration data, and the lower shows integrated corrected peak areas of the titration data fitted using the “One binding site” model. The derived dissociation constant is shown in the upper panel. Further experimental details are provided in Supplementary Table 5. **d**, **e**, Computational docking analysis of the Apo-A4_R116A, T145A, N178A, N180G mutant with epinephrine. The overall structure of the Apo-A4_R116A, T145A, N178A, N180G mutant LBD (generated by PyMOL) with epinephrine is illustrated in panel (d), with a magnified view of the ligand binding site provided in panel (e). Key residues predicted to interact with epinephrine are depicted in stick mode. Predicted interactions are depicted in dashed lines: orange for π-π stacking and black for hydrogen bonds. The mutated residues are highlighted in red.

**Extended Data Fig. 9.**
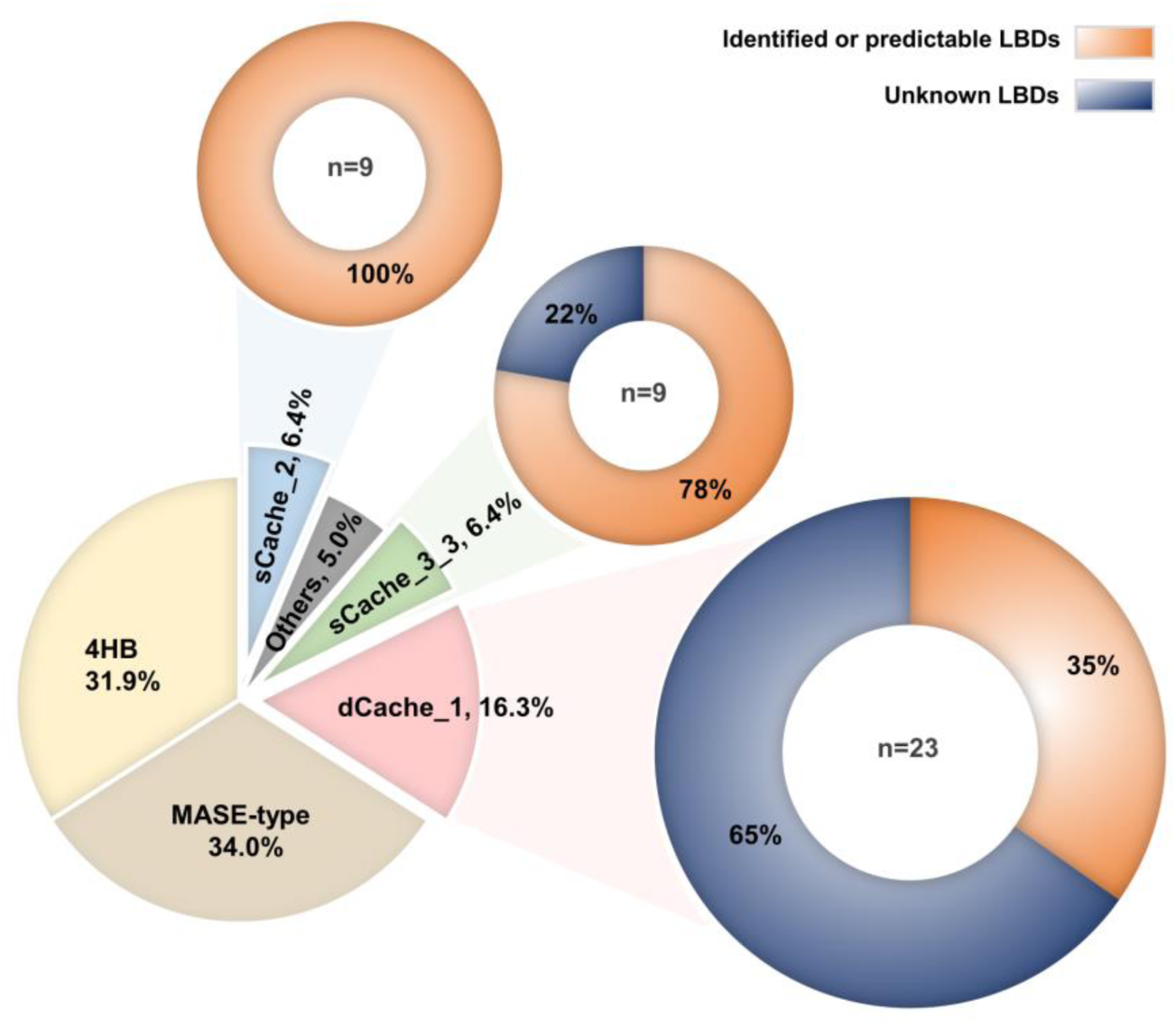
Overview of MCP specificity based on experimental identification and bioinformatic predictions. The 12 gut bacterial species analyzed possess 141 transmembrane MCPs with assigned LBDs, with the full repertoire of MCPs presented in Supplementary Data 2. Approximately one-third of these LBDs belong to the 4HB superfamily, another third to the MASE-type, and the remaining third to the Cache superfamily, which is further divided into four subgroups. Ligands of LBDs were identified experimentally or predicted using bioinformatics. All sCache_2 domains bind short-chain carboxylic acids. For sCache_3_3 domains, 78% have ligands identified through experiments or predictions based on the known formate sensors. For dCache_1 domains, 35% have ligands identified experimentally or through conserved binding motifs for amino acids, amines, or purines. Supporting details are available in Supplementary Data 2 and Extended Data Fig. 4.

**Extended Data Fig. 10.**
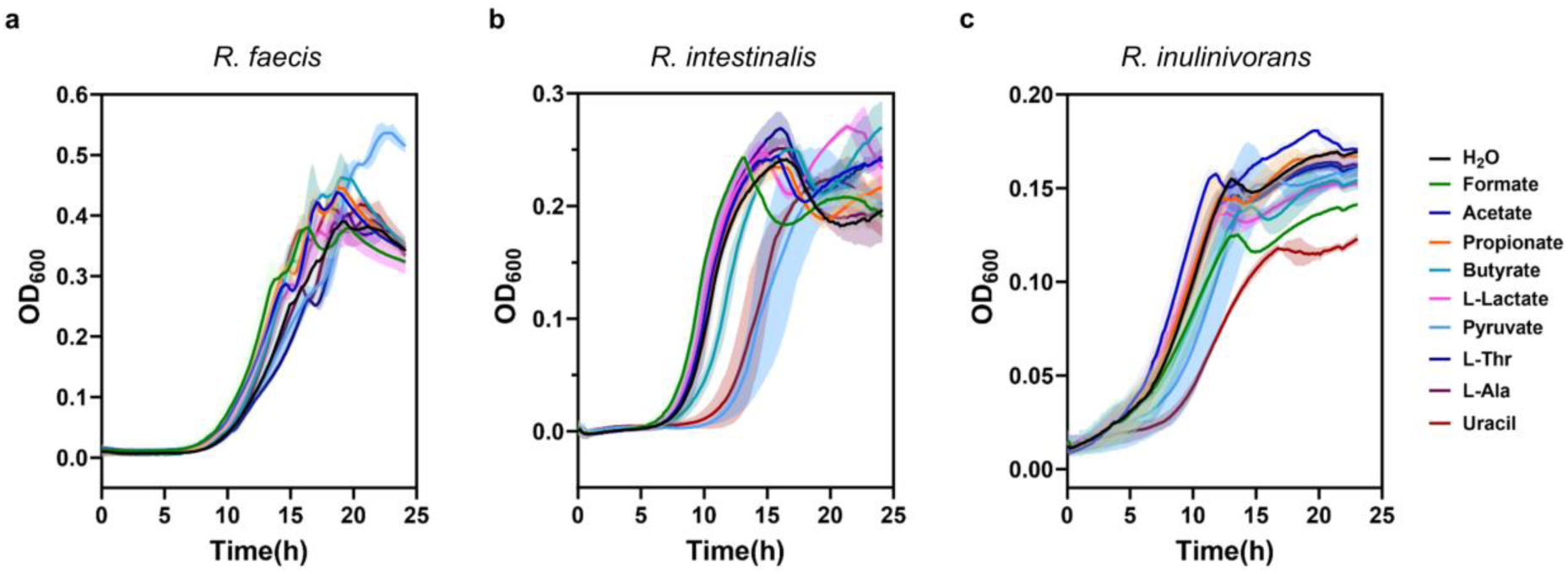
**Effects of chemoeffector compounds on growth of selected *Roseburia* species. a-c**, Growth curves of *R. faecis* (a), *R. intestinalis* (b), and *R. inulinivorans* (c) in presence of 20 mM of indicated compounds (10 mM for uracil) added to 50% YCFA medium (see Methds). Growth was assessed by measuring optical density (OD) at 600 nm (OD_600_). Solid lines represent the mean values from three biological replicates, with shaded areas indicating the standard deviations. Data represent mean values from three biological replicates

## Notes

### Competing Interest Statement

The authors have declared no competing interest.

## References

1. Fan, Y. & Pedersen, O. Gut microbiota in human metabolic health and disease. Nat. Rev. Microbiol. 19, 55–71 (2021).

2. Sommer, F., Anderson, J.M., Bharti, R., Raes, J. & Rosenstiel, P. The resilience of the intestinal microbiota influences health and disease. Nat. Rev. Microbiol. 15, 630–638 (2017).

3. Giri, S., Shi, H., Typas, A. & Huang, K.C. Harnessing gut microbial communities to unravel microbiome functions. Curr. Opin. Microbiol. 83, 102578 (2025).

4. Schofield, W.B., Zimmermann-Kogadeeva, M., Zimmermann, M., Barry, N.A. & Goodman, A.L. The stringent response determines the ability of a commensal bacterium to survive starvation and to persist in the gut. Cell Host Microbe 24, 120–132 (2018).

5. Thompson, J.A., Oliveira, R.A., Djukovic, A., Ubeda, C. & Xavier, K.B. Manipulation of the quorum sensing signal AI-2 affects the antibiotic-treated gut microbiota. Cell Rep. 10, 1861–1871 (2015).

6. Galperin, M.Y. What bacteria want. Environ. Microbiol. 20, 4221–4229 (2018).

7. Ortega, Á., Zhulin, I.B. & Krell, T. Sensory repertoire of bacterial chemoreceptors. Microbiol. Mol. Biol. Rev. 81(2017).

8. Parkinson, J.S., Hazelbauer, G.L. & Falke, J.J. Signaling and sensory adaptation in *Escherichia coli* chemoreceptors. Trends Microbiol. 23, 257–266 (2015).

9. Stock, A.M., Robinson, V.L. & Goudreau, P.N. Two-component signal transduction. Annu. Rev. Biochem. 69, 183–215 (2000).

10. Jenal, U., Reinders, A. & Lori, C. Cyclic di-GMP: second messenger extraordinaire. Nat. Rev. Microbiol. 15, 271–284 (2017).

11. Matilla, M.A., Monteagudo-Cascales, E. & Krell, T. Advances in the identification of signals and novel sensing mechanisms for signal transduction systems. Environ. Microbiol. 25, 79–86 (2023).

12. Bi, S. & Sourjik, V. Stimulus sensing and signal processing in bacterial chemotaxis. Curr. Opin. Microbiol. 45, 22–29 (2018).

13. Zhulin, I.B., Nikolskaya, A.N. & Galperin, M.Y. Common extracellular sensory domains in transmembrane receptors for diverse signal transduction pathways in bacteria and archaea. J. Bacteriol. 185, 285–294 (2003).

14. Gavira, J.A., et al. How bacterial chemoreceptors evolve novel ligand specificities. mBio 11(2020).

15. Matilla, M.A., et al. Chemotaxis of the human pathogen *Pseudomonas aeruginosa* to the neurotransmitter acetylcholine. mBio 13, e0345821 (2022).

16. Yu, Z., et al. Gas and light: triggers of c-di-GMP-mediated regulation. FEMS Microbiol. Rev. 47(2023).

17. Colin, R., Ni, B., Laganenka, L. & Sourjik, V. Multiple functions of flagellar motility and chemotaxis in bacterial physiology. FEMS Microbiol. Rev. 45(2021).

18. Keegstra, J.M., Carrara, F. & Stocker, R. The ecological roles of bacterial chemotaxis. Nat. Rev. Microbiol. 20, 491–504 (2022).

19. Nicolas, G.R. & Chang, P.V. Deciphering the chemical lexicon of host-gut microbiota interactions. Trends Pharmacol. Sci. 40, 430–430 (2019).

20. Pacheco, A.R. & Sperandio, V. Inter-kingdom signaling: chemical language between bacteria and host. Curr. Opin. Microbiol. 12, 192–198 (2009).

21. Ross, P.A., et al. Framework for exploring the sensory repertoire of the human gut microbiota. mBio 15, e0103924 (2024).

22. Lopetuso, L.R., Scaldaferri, F., Petito, V. & Gasbarrini, A. Commensal Clostridia: leading players in the maintenance of gut homeostasis. Gut Pathog. 5(2013).

23. Fernandez, M., et al. High-throughput screening to identify chemoreceptor ligands. Methods Mol. Biol. 1729, 291–301 (2018).

24. Monteagudo-Cascales, E., et al. Bacterial sensor evolved by decreasing complexity. Proc. Natl Acad. Sci. USA 122, e2409881122 (2025).

25. Sourjik, V., Vaknin, A., Shimizu, T.S. & Berg, H.C. In vivo measurement by FRET of pathway activity in bacterial chemotaxis. Methods Enzymol. 423, 365–365 (2007).

26. Xu, W., et al. Systematic mapping of chemoreceptor specificities for *Pseudomonas aeruginosa*. mBio 14, e0209923 (2023).

27. Berman, H.M., et al. The protein data bank. Acta Crystallogr. D Biol. Crystallogr. 58(2002).

28. Brewster, J.L., et al. Structural basis for ligand recognition by a Cache chemosensory domain that mediates carboxylate sensing in *Pseudomonas syringae*. Sci. Rep. 6(2016).

29. Pokkuluri, P.R., et al. Analysis of periplasmic sensor domains from *Anaeromyxobacter dehalogenans* 2CP-C: Structure of one sensor domain from a histidine kinase and another from a chemotaxis protein. MicrobiologyOpen 2(2013).

30. Remund, B., Yilmaz, B. & Sokollik, C. D-Lactate: Implications for gastrointestinal diseases. Children (Basel) 10(2023).

31. Cerna-Vargas, J.P., Gumerov, V.M., Krell, T. & Zhulin, I.B. Amine-recognizing domain in diverse receptors from bacteria and archaea evolved from the universal amino acid sensor. Proc. Natl Acad. Sci. USA 120, e2305837120 (2023).

32. Gumerov, V.M., et al. Amino acid sensor conserved from bacteria to humans. Proc. Natl Acad. Sci. USA 119, e2110415119 (2022).

33. Monteagudo-Cascales, E., et al. Ubiquitous purine sensor modulates diverse signal transduction pathways in bacteria. Nat. Commun. 15, 5867 (2024).

34. Thomas, D.M. & Zalcberg, J.R. 5-fluorouracil: a pharmacological paradigm in the use of cytotoxics. Clin. Exp. Pharmacol. Physiol. 25, 887–895 (1998).

35. Gumerov, V.M., Ulrich, L.E. & Zhulin, I.B. MiST 4.0: a new release of the microbial signal transduction database, now with a metagenomic component. Nucleic Acids Res. 52, D647–D653 (2024).

36. Nikolskaya, A.N., Mulkidjanian, A.Y., Beech, I.B. & Galperin, M.Y. MASE1 and MASE2: two novel integral membrane sensory domains. J. Mol. Microbiol. Biotechnol. 5, 11–16 (2003).

37. Matilla, M.A., Gavira, J.A. & Krell, T. Accessing nutrients as the primary benefit arising from chemotaxis. Curr. Opin. Microbiol. 75, 102358–102358 (2023).

38. Liou, M.J., et al. Host cells subdivide nutrient niches into discrete biogeographical microhabitats for gut microbes. Cell Host Microbe 30, 836–847 (2022).

39. Tropini, C., Earle, K.A., Huang, K.C. & Sonnenburg, J.L. The gut microbiome: connecting spatial organization to function. Cell Host Microbe 21, 433–442 (2017).

40. Matilla, M.A., Velando, F., Martin-Mora, D., Monteagudo-Cascales, E. & Krell, T. A catalogue of signal molecules that interact with sensor kinases, chemoreceptors and transcriptional regulators. FEMS Microbiol. Rev. 46(2022).

41. Adler, J., Hazelbauer, G.L. & Dahl, M.M. Chemotaxis toward sugars in *Escherichia coli*. J. Bacteriol. 115, 824–847 (1973).

42. Mesibov, R. & Adler, J. Chemotaxis toward amino acids in *Escherichia coli*. J. Bacteriol. 112, 315–326 (1972).

43. Johnson, K.S. & Ottemann, K.M. Colonization, localization, and inflammation: the roles of *H. pylori* chemotaxis in vivo. Curr. Opin. Microbiol. 41, 51–57 (2018).

44. Brunet, M., et al. An atlas of metabolites driving chemotaxis in prokaryotes. Nat. Commun. 16, 1242 (2025).

45. Cardona, S.T., Choy, M. & Hogan, A.M. Essential two-component systems regulating cell envelope functions: opportunities for novel antibiotic therapies. J. Membr. Biol. 251, 75–89 (2018).

46. Hirakawa, H., Inazumi, Y., Masaki, T., Hirata, T. & Yamaguchi, A. Indole induces the expression of multidrug exporter genes in *Escherichia coli*. Mol. Microbiol. 55, 1113–1126 (2005).

47. Kumar, A. & Sperandio, V. Indole signaling at the host-microbiota-pathogen interface. mBio 10(2019).

48. Tennoune, N., Andriamihaja, M. & Blachier, F. Production of indole and indole-related compounds by the intestinal microbiota and consequences for the host: the good, the bad, and the ugly. Microorganisms 10(2022).

49. Leblanc, S.K.D., Oates, C.W. & Raivio, T.L. Characterization of the induction and cellular role of the BaeSR two-component envelope stress response of *Escherichia coli*. J. Bacteriol. 193, 3367–3375 (2011).

50. Gavira, J.A., Matilla, M.A., Fernandez, M. & Krell, T. The structural basis for signal promiscuity in a bacterial chemoreceptor. FEBS J. 288, 2294–2310 (2021).

51. Guo, L., et al. Attractant and repellent induce opposing changes in the four-helix bundle ligand-binding domain of a bacterial chemoreceptor. PLoS Biol. 21, e3002429 (2023).

52. Machuca, M.A., et al. *Helicobacter pylori* chemoreceptor TlpC mediates chemotaxis to lactate. Sci. Rep. 7, 14089 (2017).

53. Chen, X., Bi, S., Ma, X., Sourjik, V. & Lai, L. Discovery of a new chemoeffector for *Escherichia coli* chemoreceptor Tsr and identification of a molecular mechanism of repellent sensing. ACS Bio and Med Chem Au 2, 386–394 (2022).

54. Matilla, M.A., Ortega, A. & Krell, T. The role of solute binding proteins in signal transduction. Comput. Struct. Biotechnol. J. 19, 1786–1805 (2021).

55. Lacal, J., García-Fontana, C., Muñoz-Martínez, F., Ramos, J.L. & Krell, T. Sensing of environmental signals: classification of chemoreceptors according to the size of their ligand binding regions. Environ. Microbiol. 12, 2873–2884 (2010).

56. Elgamoudi, B.A., et al. The *Campylobacter jejuni* chemoreceptor Tlp10 has a bimodal ligand-binding domain and specificity for multiple classes of chemoeffectors. Sci. Signal. 14(2021).

57. Pineda-Molina, E., et al. Evidence for chemoreceptors with bimodular ligand-binding regions harboring two signal-binding sites. Proc. Natl Acad. Sci. USA 109, 18926–18931 (2012).

58. Feng, H., et al. Signal binding at both modules of its dCache domain enables the McpA chemoreceptor of *Bacillus velezensis* to sense different ligands. Proc. Natl Acad. Sci. USA 119, e2201747119 (2022).

59. Mittal, R., et al. Neurotransmitters: the critical modulators regulating gut-brain axis. J. Cell. Physiol. 232, 2359–2372 (2017).

60. Louis, P., Duncan, S.H., Sheridan, P.O., Walker, A.W. & Flint, H.J. Microbial lactate utilisation and the stability of the gut microbiome. Gut Microbiome 3, e3–e3 (2022).

61. Pohanka, M. D-Lactic acid as a metabolite: toxicology, diagnosis, and detection. Biomed Res. Int., 3419034 (2020).

62. Lu, W., et al. The formate channel FocA exports the products of mixed-acid fermentation. Proc. Natl Acad. Sci. USA 109, 13254–13259 (2012).

63. Belenguer, A., et al. Two routes of metabolic cross-feeding between *Bifidobacterium adolescentis* and butyrate-producing anaerobes from the human gut. Appl. Environ. Microbiol. 72, 3593–3599 (2006).

64. De Vuyst, L. & Leroy, F. Cross-feeding between bifidobacteria and butyrate-producing colon bacteria explains bifdobacterial competitiveness, butyrate production, and gas production. Int. J. Food Microbiol. 149, 73–80 (2011).

65. Winter, M., et al. Formate oxidation in the intestinal mucus layer enhances fitness of *Salmonella enterica* serovar Typhimurium. mBio 14(2023).

66. Lopes, J.G. & Sourjik, V. Chemotaxis of *Escherichia coli* to major hormones and polyamines present in human gut. ISME J. 12, 2736–2747 (2018).

67. Hu, S. & Ottemann, K.M. *Helicobacter pylori* initiates successful gastric colonization by utilizing L-lactate to promote complement resistance. Nat. Commun. 14, 1695 (2023).

68. Li, W., et al. The EMBL-EBI bioinformatics web and programmatic tools framework. Nucleic Acids Res. 43, W580–584 (2015).

69. Zulkower, V. & Rosser, S. DNA Chisel, a versatile sequence optimizer. Bioinformatics 36, 4508–4509 (2020).

70. Gibson, D.G., et al. Enzymatic assembly of DNA molecules up to several hundred kilobases. Nat. Methods 6, 343–345 (2009).

71. Bi, S., Pollard, A.M., Yang, Y., Jin, F. & Sourjik, V. Engineering hybrid chemotaxis receptors in bacteria. ACS Synth. Biol. 5, 989–1001 (2016).

72. Velazquez-Campoy, A., Goñi, G., Peregrina, J.R. & Medina, M. Exact analysis of heterotropic interactions in proteins: Characterization of cooperative ligand binding by isothermal titration calorimetry. Biophys. J. 91(2006).

73. Paulick, A. & Sourjik, V. FRET analysis of the chemotaxis pathway response. Methods Mol. Biol. 1729, 107–126 (2018).

74. Abramson, J., et al. Accurate structure prediction of biomolecular interactions with AlphaFold 3. Nature 630, 493–500 (2024).

75. Meng, E.C., et al. UCSF ChimeraX: Tools for structure building and analysis. Protein Sci. 32, e4792 (2023).

76. Kim, S., et al. PubChem 2023 update. Nucleic Acids Res. 51, D1373–D1380 (2023).

77. Corso, G., Stärk, H., Jing, B., Barzilay, R. & Jaakkola, T. DiffDock: Diffusion steps, twists, and turns for molecular docking. Preprint at 10.48550/arXiv.2210.01776. (2022).

78. Katoh, K. & Standley, D.M. MAFFT multiple sequence alignment software version 7: Improvements in performance and usability. Mol. Biol. Evol. 30(2013).

79. Waterhouse, A.M., Procter, J.B., Martin, D.M.A., Clamp, M. & Barton, G.J. Jalview Version 2-A multiple sequence alignment editor and analysis workbench. Bioinformatics 25(2009).

80. Koblitz, J., et al. MediaDive: the expert-curated cultivation media database. Nucleic Acids Res. 51, D1531–D1538 (2023).

81. Wirth, N.T., Funk, J., Donati, S. & Nikel, P.I. QurvE: user-friendly software for the analysis of biological growth and fluorescence data. Nat. Protoc. 18, 2401–2403 (2023).

82. Kabsch, W. Integration, scaling, space-group assignment and post-refinement. Acta Crystallogr. D Biol. Crystallogr. 66, 133–144 (2010).

83. McCoy, A.J., et al. Phaser crystallographic software. J. Appl. Crystallogr. 40, 658–674 (2007).

84. Emsley, P., Lohkamp, B., Scott, W.G. & Cowtan, K. Features and development of Coot. Acta Crystallogr. D Biol. Crystallogr. 66, 486–501 (2010).

85. Liebschner, D., et al. Macromolecular structure determination using X-rays, neutrons and electrons: recent developments in Phenix. Acta Crystallogr. D Struct. Biol. 75, 861–877 (2019).

86. Laskowski, R.A., Jablonska, J., Pravda, L., Varekova, R.S. & Thornton, J.M. PDBsum: Structural summaries of PDB entries. Protein Sci. 27, 129–134 (2018).

87. Capella-Gutierrez, S., Silla-Martinez, J.M. & Gabaldon, T. trimAl: a tool for automated alignment trimming in large-scale phylogenetic analyses. Bioinformatics 25, 1972–1973 (2009).

88. Darriba, D., Taboada, G.L., Doallo, R. & Posada, D. ProtTest 3: fast selection of best-fit models of protein evolution. Bioinformatics 27, 1164–1165 (2011).

89. Ronquist, F., et al. MrBayes 3.2: efficient Bayesian phylogenetic inference and model choice across a large model space. Syst. Biol. 61, 539–542 (2012).

