## Supplementary Information for "Specificities of Chemosensory Receptors in the Human Gut Microbiota"

to

by

Wenhao Xu, Ekaterina Jalomo-Khayrova, Vadim M Gumerov, Patricia A. Ross, Tania S. Köbe, Daniel Schindler, Gert Bange, Igor B. Zhulin, Victor Sourjik

**Table of content**

- Supplementary Figures 1-3
- Supplementary Tables 1-6
- Supplementary References

### Supplementary Figures

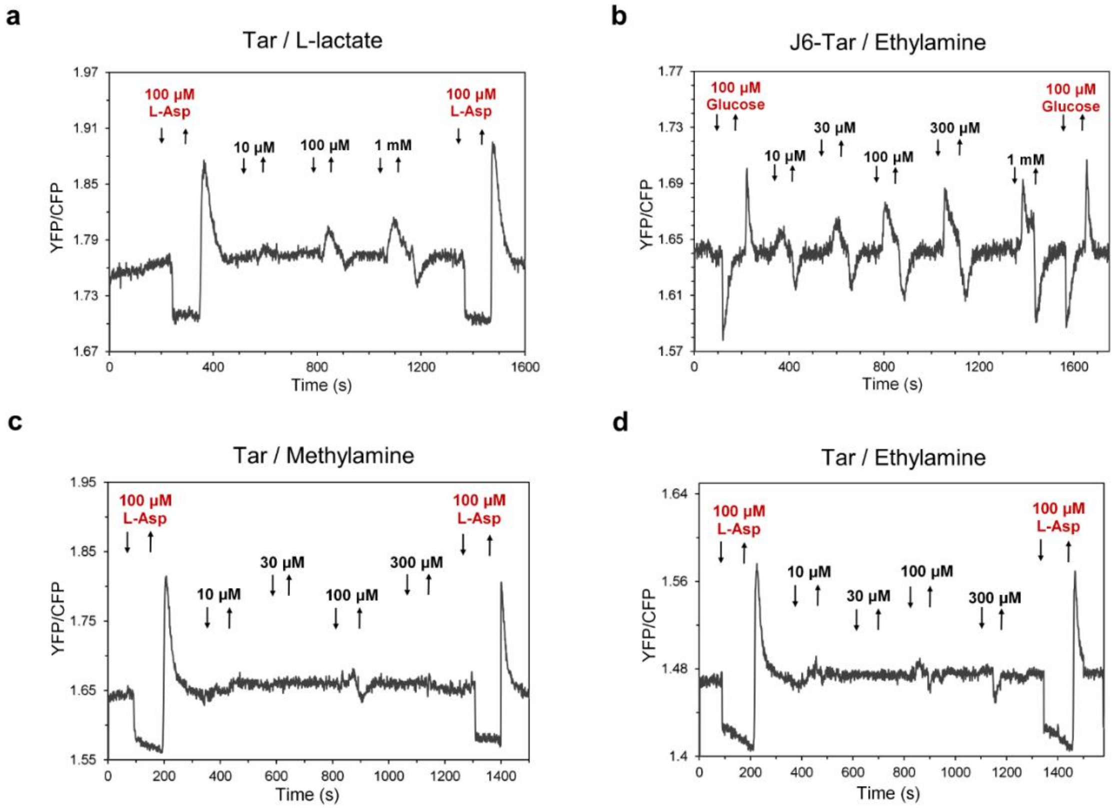


**Supplementary Fig. 1 Confirmation of signal transduction mediated by the K1-Tar and J6-Tar chimeras using FRET. a**, FRET measurement of the wild-type Tar response to L-lactate. Buffer-adapted *E. coli* cells expressing the CheZ-CFP/CheY-YFP FRET pair and Tar as the sole receptor stimulated by stepwise addition (down arrow) and subsequent removal (up arrow) of the indicated concentrations of L-lactate. Tar ligand L-asparate was used as a positive control. **b**, FRET measurement of the hybrid chemoreceptor J6-Tar response to ethylamine. D-glucose was used as a positive control ligand to confirm the activity of chimera. **c**, **d**, FRET measurement of the wild-type Tar response to methylamine (c) or ethylamine (d).

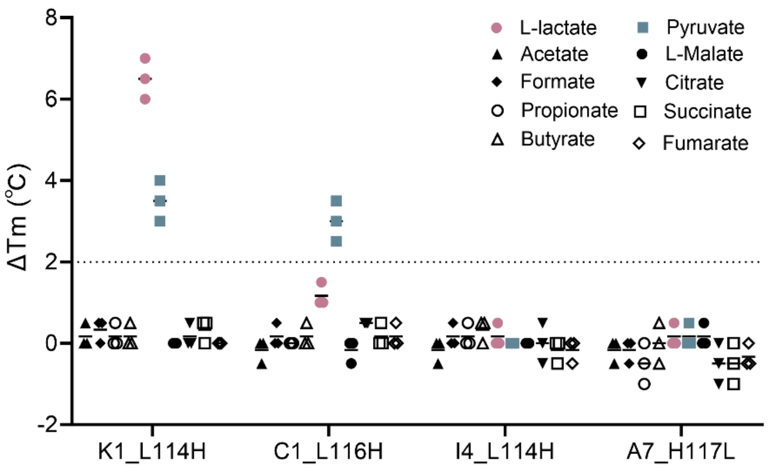


**Supplementary Fig. 2 Binding studies of four mutated sCache_2 SCCA domains.** Theram shift measurements for sensor proteins mutants with 2 mM concentrations of indicated short-chain carboxylic acids (SCCAs). Each data point represents the individual biological measurement. Short black lines represent the mean of three independent biological replicates. The black dashed line indicates the threshold of 2 °C used as a significance cutoff for ligand identification.

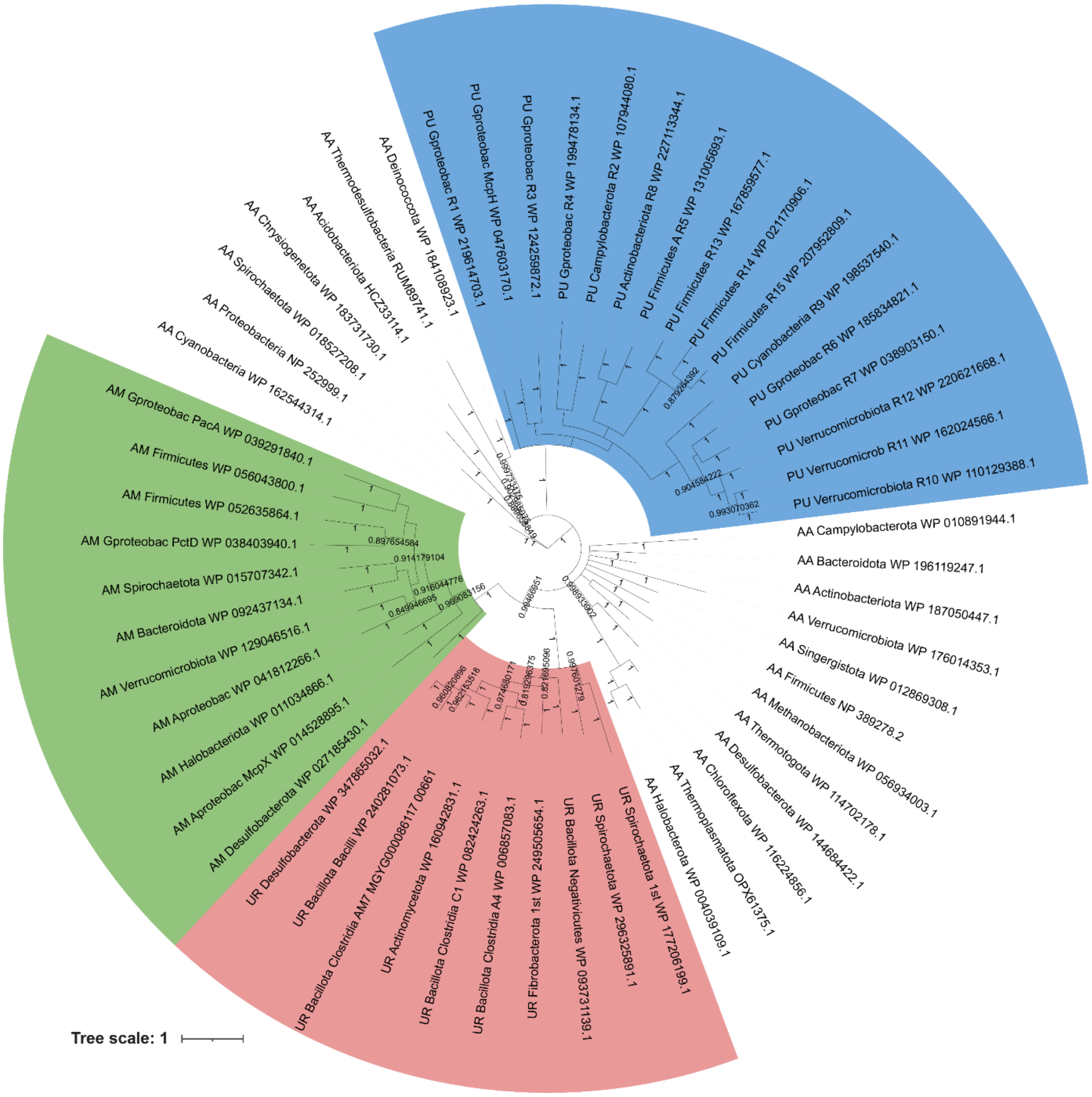


**Supplementary Fig. 3 Bayesian phylogenetic tree depicted in Fig. 5b with annotation of individual proteins.** Colors and abbreviations are as in Fig. 5b.

### Supplementary Tables

**Supplementary Table 1. Source gut bacteria of the studied sensory domains.**

| NCBI Organism name[1] | GTDB Taxonomy | Genome Accession | No.[2] |
|---|---|---|---|
| *Agathobacter rectalis* | d__Bacteria; p__Bacillota_A; c__Clostridia; o__Lachnospirales; f__Lachnospiraceae; g__Agathobacter; s__Agathobacter rectalis | GCF_001404855.1 | 6 |
| *[Eubacterium] siraeum* DSM 15702 | d__Bacteria; p__Bacillota_A; c__Clostridia; o__Oscillospirales; f__Ruminococcaceae; g__Ruminiclostridium_E; s__Ruminiclostridium_E siraeum | GCF_000382085.1 | 4 |
| *Blautia obeum* ATCC 29174 | d__Bacteria; p__Bacillota_A; c__Clostridia; o__Lachnospirales; f__Lachnospiraceae; g__Blautia_A; s__Blautia_A obeum | GCF_000153905.1 | 2 |
| *Blautia wexlerae* DSM 19850 | d__Bacteria; p__Bacillota_A; c__Clostridia; o__Lachnospirales; f__Lachnospiraceae; g__Blautia_A; s__Blautia_A wexlerae | GCF_000484655.1 | 17 |
| *Catenibacterium mitsuokai* DSM 15897 | d__Bacteria; p__Bacillota_I; c__Bacilli_A; o__Erysipelotrichales; f__Coprobacillaceae; g__Catenibacterium; s__Catenibacterium mitsuokai | GCF_000173795.1 | 3 |
| *Collinsella aerofaciens* | d__Bacteria; p__Actinomycetota; c__Coriobacteriia; o__Coriobacteriales; f__Coriobacteriaceae; g__Collinsella; s__Collinsella aerofaciens_A | GCF_002736145.1 | 1 |
| *Eisenbergiella massiliensis* | d__Bacteria; p__Bacillota_A; c__Clostridia; o__Lachnospirales; f__Lachnospiraceae; g__Eisenbergiella; s__Eisenbergiella porci | GCF_900243045.1 | 6 |
| *Lachnospira eligens* | d__Bacteria; p__Bacillota_A; c__Clostridia; o__Lachnospirales; f__Lachnospiraceae; g__Lachnospira; s__Lachnospira eligens_A | GCF_001405395.1 | 6 |
| *Fusicatenibacter saccharivorans* | d__Bacteria; p__Bacillota_A; c__Clostridia; o__Lachnospirales; f__Lachnospiraceae; g__Fusicatenibacter; s__Fusicatenibacter saccharivorans | GCF_001406335.1 | 1 |
| *Hungatella hathewayi* | d__Bacteria; p__Bacillota_A; c__Clostridia; o__Lachnospirales; f__Lachnospiraceae; g__Hungatella; s__ Hungatella effluvii | GCF_001405675.1 | 1 |
| *Lachnospira pectinoschiza* | d__Bacteria; p__Bacillota_A; c__Clostridia; o__Lachnospirales; f__Lachnospiraceae; g__Lachnospira; s__Lachnospira pectinoschiza_A | GCF_001405835.1 | 15 |
| *Lactobacillus rogosae* | d__Bacteria; p__Bacillota_A; c__Clostridia; o__Lachnospirales; f__Lachnospiraceae; g__Lachnospira; s__Lachnospira pectinoschiza_A | GCF_900112995.1 | 5 |
| *Phascolarctobacterium faecium* DSM 14760 | d__Bacteria; p__Bacillota_C; c__Negativicutes; o__Acidaminococcales; f__Acidaminococcaceae; g__Phascolarctobacterium; s__Phascolarctobacterium faecium | GCF_003269275.1 | 1 |
| *Roseburia faecis* | d__Bacteria; p__Bacillota_A; c__Clostridia; o__Lachnospirales; f__Lachnospiraceae; g__Agathobacter; s__Agathobacter faecis | GCF_001405615.1 | 19 |
| *Roseburia hominis* A2-183 | d__Bacteria; p__Bacillota_A; c__Clostridia; o__Lachnospirales; f__Lachnospiraceae; g__Roseburia; s__Roseburia hominis | GCF_000225345.1 | 6 |
| *Roseburia intestinalis* L1-82 | d d__Bacteria; p__Bacillota_A; c__Clostridia; o__Lachnospirales; f__Lachnospiraceae; g__Roseburia; s__Roseburia intestinalis | GCF_000156535.1 | 11 |
| *Roseburia inulinivorans* DSM 16841 | d__Bacteria; p__Bacillota_A; c__Clostridia; o__Lachnospirales; f__Lachnospiraceae; g__Roseburia; s__Roseburia inulinivorans | GCF_000174195.1 | 8 |
| *Ruminococcus bicirculans* | d__Bacteria; p__Bacillota_A; c__Clostridia; o__Oscillospirales; f__Ruminococcaceae; g__Hominimerdicola; s__Hominimerdicola aceti | GCF_000723465.1 | 1 |
| *Ruminococcus bromii* | d__Bacteria; p__Bacillota_A; c__Clostridia; o__Oscillospirales; f__Acutalibacteraceae; g__Ruminococcus_E; s__Ruminococcus_E bromii_A | GCF_900101355.1 | 1 |
| *Subdoligranulum* sp. 4_3_54A2FAA | d__Bacteria; p__Bacillota_A; c__Clostridia; o__Oscillospirales; f__Ruminococcaceae; g__Ruthenibacterium; s__Ruthenibacterium lactatiformans | GCF_000238635.1 | 2 |

[1]The NCBI organism name was used throughout this study to refer to the related species.
[2]The number of sensor proteins studied from the specified organism is indicated.

**Supplementary Table 2. Layouts of the human gut metabolite plates (HGMT plates).** All compounds were dissolved in $H_2O$ at the final concentration of 20 mM and pH 7.0, unless otherwise stated. Different chemical categories are highlighted with color.

**HGMT Plate_1**

| | A | B | C | D | E | F | G | H |
|---|---|---|---|---|---|---|---|---|
| **1** | $H_2O$ | L-Lysine | D-Aspartic acid | Ethanolamine | Acetylcholine | Pyruvate | 4-Methylaminobutyric acid | D-Quinic acid |
| **2** | L-Alanine | L-Methionine | D-Serine | L-Carnitine | Choline | L-Malate | Glutathione reduced | Ferulic acid |
| **3** | L-Arginine | L-Phenylalanine | Ala-Gln | Betaine | Isobutyric acid | Fumarate | 3-Hydroxypropionic acid | Trans-Aconitic acid |
| **4** | L-Asparagine | L-Proline | Gly-Glu | Spermidine[a] | Formate | Tartrate | β-Hydroxybutyric acid | α-Ketoglutaric acid |
| **5** | L-Aspartic acid | L-Serine | N-Acetyl-L-Glutamate (GluNAC) | Spermine[a] | Acetate | Oxalacetic acid | 5-Aminovalerate | |
| **6** | L-Cysteine | L-Threonine | N-Acetyl-L-aspartic acid | Agmatine | Propionate | Glutaric acid | Methylmalonic acid | |
| **7** | L-Glutamine | L-Tryptophan | Gly-Gly-Ala | Putrescine | Isovaleric acid | L-Lactate | Tricarballylate | |
| **8** | L-Glutamic acid | L-Tyrosine (KOH) | Creatine Monohydrate | Phenethylamine | Butyrate | Succinate | α-Aminobutyric acid | |
| **9** | L-Glycine | L-Valine | Dimethylamine hydrochloride | Histamine | Citrate | Phenylacetate | Glyoxylic acid monohydrate | |
| **10** | L-Histidine | L-Ornithine | Methylamine hydrochloride | Trimethylamine | Gluconic acid | Salicylate | γ-aminobutyric acid | |
| **11** | L-Isoleucine | L-Citrulline | Tyramine hydrochloride | Ethylenediamine | Malonate | Caffeic acid | Maleate | |
| **12** | L-Leucine | D-Alanine | Trimethylamine N-oxide (TMAO) | Ethylamine | Oxalic acid | Itaconate | Shikimic acid | |

**HGMT Plate_2**

| | A | B | C | D | E | F |
|---|---|---|---|---|---|---|
| 1 | $H_2O$ | L-Fucose | D-Trehalose | Sodium cholate[b] | Hypoxanthine (10 mM) | Inosine |
| 2 | D-Xylose | α-D-Raffinose • $H_2O$ | Inulin | Taurine | Adenosine | Uridine |
| 3 | D-Mannitol | D-Galactose | L-Arabinose | Urea | Purine | Cytosine (10 mM) |
| 4 | D-Fructose | Glucosamine | Ascorbate | Acetamide | Uric acid (50 mM KOH) | Theophylline |
| 5 | D-Maltose | Methyl-α-D-Glucopyranoside | Biotin | $NaNO_2$ | Xanthine (50 mM KOH ) | Thymidine |
| 6 | Lactose | D-Glucose-6-phosphate | Vitamin B1 (Thiamin) | $NaNO_3$ | Guanine (50 mM KOH ) | |
| 7 | D-Mannose | D-Ribose | Myo-inositol | Indole (10 mM) | Guanosine (50 mM KOH ) | |
| 8 | D-Glucose | N-Acetyl-D-Glucosamine (GlcNAC)[a] | Nicotinic acid | Allantoin | Adenine (10 mM) | |
| 9 | D-Sucrose | N-Acetyl-D-Galactose (GalNAC) | Nicotinamide | KOH (50 mM) | Uracil (10 mM) | |
| 10 | Xylan | D-Fructose-6-phosphate[a] | Pyridoxine hydrochloride | (±)-Epinephrine hydrochloride | Thymine (10 mM) | |
| 11 | D-Sorbitol | D-Ribose-5-phosphate[a] | Hyocholic acid (10 mM) | Dopamine hydrochloride | Cytidine | |
| 12 | L-Rhamnose | Isomaltulose | Sodium taurocholate hydrate[b] | 3,4-Dihydroxymandelic acid | Caffeine | |

| | | | | |
|---|---|---|---|---|
| Amino acids and derivatives | Biogenic amines | Short-chain fatty acids | Other carboxylic acids | Sugars and derivatives |
| Vitamins | Bile acids | Hormones | Nucleobases and derivatives | Other compounds |

[a] Compounds that caused thermal shifts in multiple ligand binding domains, possibly because these ligands induce nonspecific conformational changes in LBDs.

[b] Compounds that caused protein unfolding in thermal shift assays.

**Supplementary Table 3. Reported concentrations of selected metabolites in the mammalian intestine.**

| Metabolite | Ligand | Sample source | Concentration | Ref. |
|---|---|---|---|---|
| Amino acids | L-arginine | Human, intestinal lumen | Free amino acid concentration: 50-300 µM, 0.6-6 mM after a protein-rich meal | 1 |
| | L-valine | | | |
| | L-threonine | | | |
| | L-glycine | | | |
| | L-alanine | | | |
| | D-serine | | | |
| Biogenic amines | Methylamine | Human, colon | Proximal colon: 0-6.0 mM; distal colon: 2.2-3.2 mM | 2 |
| | Ethylamine | - | - | - |
| Nucleobases and their derivatives | Inosine | Mice, GI tract | Duodenum: 66.13±14.23 µM; jejunum: 29.26±9.38 µM; cecum 0.5±0.05 µM | 3 |
| | Hypoxanthine | Mice, small intestine | 0-0.3 mM | 4 |
| | Theophylline | Rabbit, GI tract | Up to10 mM (depending on diet) | 5 |
| | Uracil | Mice, small intestinal | 0-0.5 mM | 4 |
| C3/C4-dicarboxylic acids | Succinate | Human, feces | 6.3±1.7 mM (in healthy control) | 6 |
| | Itaconate | - | - | |
| | Maleate | - | - | |
| | Methymalonate | - | - | |
| Short-chain fatty acids | Formate | Human, colon | 0-5mM | 7 |
| | Acetate | Human, feces | 17.9-164.1 mM; average: 62.4 mM | 8 |
| | Propionate | Human, feces | 4.3-49.8 mM; average: 21.0 mM | 8 |
| | Butyrate | Human, feces | 1.6-70.1 mM; average: 18.8 mM | 8 |
| Other short-chain carboxylic acids | Pyruvate | - | - | - |
| | L-lactate | Human, feces | 0-25 mM | 9 |
| Sugar | D-fructose | Human, small intestine | 6-15 mM (under fasting conditions) | 10 |
| Indole | Indole | Human, GI tract | 0.2-6.5 mM | 11 |

**Supplementary Table 4. Strains, plasmids, and oligonucleotides used in this study.**

| Strains and plasmids | Genotype or relevant characteristics[a] | Reference |
|---|---|---|
| Strains | | |
| T7 Express (Enhanced *E. coli* BL21 derivative) | *fhuA2 lacZ::T7 gene1 [lon] ompT gal sulA11 R(mcr-73::miniTn10*--Tet$^{S}$*)2 [dcm] R(zgb-210::Tn10*--Tet$^{S}$*) endA1 Δ(mcrC-mrr)114::IS10* | New England Biolabs, [12] |
| *E. coli* DH5α | F$^{–}$ φ80*lac*ZΔM15 Δ(*lac*ZYA-*arg*F) U169 *rec*A1 *end*A1 *hsd*R17(r$_{K}^{–}$, m$_{K}^{+}$) *pho*A *sup*E44 λ$^{–}$*thi*-1 *gyr*A96 *rel*A1 | [13] |
| *E. coli* UU1250 | Derivative of RP437; Δ*aer*Δ*tsr*Δ(*tar-tap*) Δ*trg* | [14] |
| *E. coli* VS181 | Derivative of RP437; Δ(*cheYcheZ*)Δ*aer*Δ*tsr*Δ(*tar-tap*) Δ*trg* | [15] |
| Plasmids | | |
| pET28a (+) | Km$^{R}$; Protein expression vector | Novagen |
| pKG116 | Cm$^{R}$; Expression vector, salicylate inducible; for generation of hybrid chemoreceptor | [16] |
| pSB13 | Cm$^{R}$; Tar expression plasmid, pKG116 derivative, T768 was mutated to C768 to remove the NdeI restriction site | [17] |
| pVS88 | Ap$^{R}$; CheY-EYFP / CheZ-ECFP expression plasmid | [15] |
| pET28_LBDs library | Km$^{R}$; pET28a (+) derivative containing a DNA fragment encoding individual full-length LBD from LBD library | This study |
| pET28_K-Y101A | Km$^{R}$; pET28a (+) derivative containing a DNA fragment encoding the K1$_{LBD}$-Y101A mutant | This study |
| pET28_K1-W103A | Km$^{R}$; pET28a (+) derivative containing a DNA fragment encoding the K1$_{LBD}$-W103A mutant | This study |
| pET28_K1-L114A | Km$^{R}$; pET28a (+) derivative containing a DNA fragment encoding the K1$_{LBD}$-L114A mutant | This study |
| pET28_K1-M135A | Km$^{R}$; pET28a (+) derivative containing a DNA fragment encoding the K1$_{LBD}$-M135A mutant | This study |
| pET28_K1-Y153A | Km$^{R}$; pET28a (+) derivative containing a DNA fragment encoding the K1$_{LBD}$-Y153A mutant | This study |
| pET28_K1-F155A | Km$^{R}$; pET28a (+) derivative containing a DNA fragment encoding the K1$_{LBD}$-F155A mutant | This study |
| pET28_K1-K166A | Km$^{R}$; pET28a (+) derivative containing a DNA fragment encoding the K1$_{LBD}$-K166A mutant | This study |
| pET28_ C1-L116H | Km$^{R}$; pET28a (+) derivative containing a DNA fragment encoding the C1$_{LBD}$-L116H mutant | This study |

| | | |
|---|---|---|
| pET28_A7-H117L | $Km^R$; pET28a (+) derivative containing a DNA fragment encoding the $A7_{LBD}$-H117L mutant | This study |
| pET28_ K1-L114H | $Km^R$; pET28a (+) derivative containing a DNA fragment encoding the $K1_{LBD}$-L114H mutant | This study |
| pET28_I4-L114H | $Km^R$; pET28a (+) derivative containing a DNA fragment encoding the $I4_{LBD}$-L114H mutant | This study |
| pET28_D8-dm | $Km^R$; pET28a (+) derivative containing a DNA fragment encoding the distal module of $D8_{LBD}$ | This study |
| pET28_D8-pm | $Km^R$; pET28a (+) derivative containing a DNA fragment encoding the proximal module of $D8_{LBD}$ | This study |
| pET28_C1-dm | $Km^R$; pET28a (+) derivative containing a DNA fragment encoding the distal module of $C1_{LBD}$ | This study |
| pET28_A4-dm | $Km^R$; pET28a (+) derivative containing a DNA fragment encoding the distal module of $A4_{LBD}$ | This study |
| pET28_M8-dm | $Km^R$; pET28a (+) derivative containing a DNA fragment encoding the distal module of $M8_{LBD}$ | This study |
| pET28_A4-R116A | $Km^R$; pET28a (+) derivative containing a DNA fragment encoding the $A4_{LBD}$-R116A mutant | This study |
| pET28_A4-F129A | $Km^R$; pET28a (+) derivative containing a DNA fragment encoding the $A4_{LBD}$-F129A mutant | This study |
| pET28_A4-T145A | $Km^R$; pET28a (+) derivative containing a DNA fragment encoding the $A4_{LBD}$-T145A mutant | This study |
| pET28_A4-W160A | $Km^R$; pET28a (+) derivative containing a DNA fragment encoding the $A4_{LBD}$-W160A mutant | This study |
| pET28_A4-Y176A | $Km^R$; pET28a (+) derivative containing a DNA fragment encoding the $A4_{LBD}$-Y176A mutant | This study |
| pET28_A4-N178A | $Km^R$; pET28a (+) derivative containing a DNA fragment encoding the $A4_{LBD}$-N178A mutant | This study |
| pET28_A4-N180A | $Km^R$; pET28a (+) derivative containing a DNA fragment encoding the $A4_{LBD}$-N180A mutant | This study |
| pET28_A4-D205A | $Km^R$; pET28a (+) derivative containing a DNA fragment encoding the $A4_{LBD}$-D205A mutant | This study |
| pET28_A4-Y225A | $Km^R$; pET28a (+) derivative containing a DNA fragment encoding the $A4_{LBD}$-Y225A mutant | This study |
| pET28_A4-H238A | $Km^R$; pET28a (+) derivative containing a DNA fragment encoding the $A4_{LBD}$-H238A mutant | This study |

| | | |
|---|---|---|
| pET28_A4-Y273A | $Km^R$; pET28a (+) derivative containing a DNA fragment encoding the $A4_{LBD}$-Y273A mutant | This study |
| pET28_A4-K280A | $Km^R$; pET28a (+) derivative containing a DNA fragment encoding the $A4_{LBD}$-K280A mutant | This study |
| pET28_A4-R116A, T145A, N178A, N180G | $Km^R$; pET28a (+) derivative containing a DNA fragment encoding the $A4_{LBD}$-R116A, T145A, N178A, N180G mutant | This study |
| J6-Tar | $Cm^R$; pKG116 derivative containing a DNA fragment encoding J6 [1-345]-SLLPY-Tar [203-553]; | This study |
| K1-Tar | $Cm^R$; pKG116 derivative containing a DNA fragment encoding K1 [1-218]-LSVRL-Tar [203-553]; | This study |
| K1-Tar-L114H | $Cm^R$; pKG116 derivative containing a DNA fragment encoding K1 [1-218]-LSVRL-Tar [203-553]; L114 was mutated to H114; | This study |

[a]Ap, ampicillin; Km, kanamycin; Tc, tetracycline; Cm, chloramphenicol; Sm, streptomycin

**Oligonucleotides**

| Oligonucleotide | Sequence (5'-3') | Purpose |
|---|---|---|
| K1_ Y101A_F | TGAGGCCGGCgcgTTCTGGGTCG | Construction of pET28_ $K1_{LBD}$_ Y101A |
| K1_ Y101A _R | CCGTAACGCATTTGG | |
| K1_ W103A _F | CGGCTATTTCgcgGTCGATCAATCCG | Construction of pET28_ $K1_{LBD}$_ W103A |
| K1_ W103A _R | GCCTCACCGTAACGC | |
| K1_ L114A _F | AAATATAGTGgcgCTCGGCTCGAG | Construction of pET28_ $K1_{LBD}$_ L114A |
| K1_ L114A _R | TTACCATCGGATTGATC | |
| K1_ M135A _F | TGGATATCAGgcgGTGAAAGAAATTATTCG | Construction of pET28_ $K1_{LBD}$_ M135A |
| K1_ M135A _R | TCGGCGTCTTTGGTG | |
| K1_ Y153A _F | CTATACAGATgcgGTTTTTCCGAAGGAAGGTGAAACCG | Construction of pET28_ $K1_{LBD}$_ Y153A |
| K1_ Y153A _R | CCCCCGCCATCCTGT | |
| K1_ F155A _F | AGATTACGTTgcgCCGAAGGAAGGTG | Construction of pET28_ $K1_{LBD}$_ F155A |
| K1_ F155A _R | GTATAGCCCCCGCCA | |
| K1_ K166A _F | ACCATCACCTgcgCGCAGTTACTC | Construction of pET28_ $K1_{LBD}$_ K166A |
| K1_ K166A _R | TCGGTTTCACCTTCC | |
| K1_ L114H _F | AAATATAGTGcatCTCGGCTCGAG | Construction of pET28_ $K1_{LBD}$_ L114H |
| K1_ L114H _R | TTACCATCGGATTGATC | |

| | | |
|---|---|---|
| C1_ L116H _F | GAACGTGGTCcatTTGGGTAATGATAC | Construction of pET28_ $C1_{LBD}$_ L116H |
| C1_ L116H _R | GTCCCATCATAGGTGTC | |
| I4_ L114H _F | GAACGTCGTTcatCTGGGTTCGG | Construction of pET28_ $I4_{LBD}$_ L114H |
| I4_ L114H _R | GTGCCATCACTCTGATC | |
| A7_ H117L_F | CCTCATCATGctgCCGATTCTGAC | Construction of pET28_ $A7_{LBD}$_ H117L |
| A7_ H117L_R | TATAATCAGTATCATCAATCCAAAAG | |
| D8-dm_F | GTGCCGCGCGGCAGCCATATGAGTACTACGAAAGCACTGACC | Construction of pET28_ $D8_{LBD}$-dm |
| D8-dm_R | ACGGAGCTCGAATTCGGATCCCTAACGCACGATATCGTTCAGAA | |
| D8-pm_F | GTGCCGCGCGGCAGCCATATGAAAAACGGTGTTTTAAATAGTGAAG | Construction of pET28_ $D8_{LBD}$-pm |
| D8-pm_R | ACGGAGCTCGAATTCGGATCCCTATTTCAGGCAATGGATGGTATAAT | |
| H8-dm_F | GTGCCGCGCGGCAGCCATATGTCCGTACGTACCGCG | Construction of pET28_ $H8_{LBD}$-dm |
| H8-dm_R | ACGGAGCTCGAATTCGGATCCCTAGGTTTCATCCACCAGATCG | |
| A4-dm_F | GTGCCGCGCGGCAGCCATATGAGTACCTCGTATGAAGATAGCC | Construction of pET28_ $A4_{LBD}$-dm |
| A4-dm_R | ACGGAGCTCGAATTCGGATCCCTATGTATCGGTGAACTCGCTA | |
| M8-dm_F | GTGCCGCGCGGCAGCCATATGAAGACCGTCGTTGATGAAG | Construction of pET28_ $M8_{LBD}$-dm |
| M8-dm_R | ACGGAGCTCGAATTCGGATCCCTATTCCCGAATAACATCATAGTTG | |
| A4_R116A_F | CGCCTACATTgcgTATAATCCAGAATTTACG | Construction of pET28_ $A4_{LBD}$_R116A |
| A4_R116A_R | GTCAGGGCACCTTTC | |
| A4_F129A_F | AAGCGGCCTGgcgCTGACCCGTG | Construction of pET28_ $A4_{LBD}$_F129A |
| A4_F129A_R | GTGGGTTCCGTAAATTCTGG | |
| A4_T145A_F | CGTTACTCCAgcgGATTTTAGCATG | Construction of pET28_ $A4_{LBD}$_T145A |
| A4_T145A_R | CTCTCAAATTCACTATCCG | |
| A4_W160A_F | ACATGTCGGGgcgTATTACATTCCTG | Construction of pET28_ $A4_{LBD}$_W160A |
| A4_W160A_R | TCTACGTCGCTCGGATC | |

| | | |
|---|---|---|
| A4_Y176A_F | GATGGAGCCTgcgCTGAACTCCAATATTG | Construction of pET28_ A4$_{LBD}$_Y176A |
| A4_Y176A_R | CAGGTTTCTTTACCATTCTG | |
| A4_N178A_F | GCCTTATCTGgcgTCCAATATTGGAGTG | Construction of pET28_ A4$_{LBD}$_N178A |
| A4_N178A_R | TCCATCCAGGTTTCTTTAC | |
| A4_N180G_F | TCTGAACTCCggcATTGGAGTGTAC | Construction of pET28_ A4$_{LBD}$_N180G |
| A4_N180G_R | TAAGGCTCCATCCAG | |
| A4_N180A_F | TCTGAACTCCgcgATTGGAGTGTAC | Construction of pET28_ A4$_{LBD}$_N180A |
| A4_N180A_R | TAAGGCTCCATCCAG | |
| A4_D205A_F | CATTGGAATGgcgATTGATTTTAGCG | Construction of pET28_ A4$_{LBD}$_D205A |
| A4_D205A_R | ATGCCGATAGATTCAC | |
| A4_Y225A_F | CGACTCTGGCgcgGGATTTCTTGTG | Construction of pET28_ A4$_{LBD}$_Y225A |
| A4_Y225A_R | AAAATGCTAAGACTATCAATTG | |
| A4_H238A_F | GGTGATGTACgcgAAAGATCTGGAAATCG | Construction of pET28_ A4$_{LBD}$_H238A |
| A4_H238A_R | TTTCCGGATTCATTCAC | |
| A4_Y273A_F | CGCGGTGAGCgcgACCTACCAGG | Construction of pET28_ A4$_{LBD}$_Y273A |
| A4_Y273A_R | GTTTCCTCAGTCTGTTC | |
| A4_K280A_F | GGGAAAGGATgcgGTGATGTATTATAAGAC | Construction of pET28_ A4$_{LBD}$_K280A |
| A4_K280A_R | TGGTAGGTGTAGCTC | |
| pKG116-seq_F | AAGCCATAAGGAGTACCATATG | Sequencing for hybrid Chemoreceptor |
| pKG116-seq_R | TTACTTATTTATCCGCGGATC | |
| pET28a-seq_F | GTGCCGCGCGGCAGCCATATG | Sequencing for LBD expression plasmids |
| pET28a-seq_R | ACGGAGCTCGAATTCGGATCCCTA | |
| J6-Tar_F | AGCCATAAGGAGTACCATATGATGAGCAAAGAAC ACACG | Construction of J6-Tar chimera |
| J6-Tar _R | CCACCAGCAGAATCAANNNNNNNNNNNNNNNNAA CAATCATAAATACCACTAGAATA | |
| K1-Tar_F | AGCCATAAGGAGTACCATATGATGAAAAATATTA AAGTCCGCACG | Construction of K1-Tar chimera |
| K1-Tar_R | CCACCAGCAGAATCAANNNNNNNNNNNNNNNNCA TACAAACGCTGCACAGG | |

**Supplementary Table 5. Microcalorimetric analysis of ligand binding to sensory domains from gut microbiota.** For each protein, its affinities, experimental conditions, and enthalpy changes induced by the addition of relevant ligands were shown.

| **Protein** | | | **Ligand** | | **$K_D$ (µM)** | **$K_D$ Error (µM)** | **ΔH (kcal/mol)** | **ΔH Error (kcal/mol)** |
|---|---|---|---|---|---|---|---|---|
| **Pfam domain family** | **Protein ID** | **Concentration (µM)** | **Name** | **Concentration (mM)** | | | | |
| SMP_2 | B6 | 69 | Indole | 2 | 35.6 | 7.02 | -6.2 | 1.5 |
| sCache_2 | K1 | 85 | L-lactate | 1 | 16.0 | 0.56 | -26.9 | 0.5 |
| | | 87 | D-lactate | 3 | 66.1 | 10.7 | -5.41 | 0.5 |
| | | 85 | D-fructose | 0.3 | N/A | N/A | N/A | N/A |
| | C1 | 69 | L-lactate | 1 | 26.8 | 0.91 | -51.0 | 1.7 |
| dCache_1 | B9 | 25 | L-threonine | 0.2 | 1.3 | 0.20 | -19.5 | 1.1 |
| | | 81 | L-valine | 1.5 | N/A | N/A | N/A | N/A |
| | J6 | 73 | Methylamine | 2 | 242.0 | 22.00 | -29.7 | 9.2 |
| | | 73 | Ethylamine | 2 | 134.0 | 8.30 | -55.1 | 16.3 |
| | D8 | 63 | Succinate | 0.5 | 1.7 | 0.11 | -21.9 | 0.3 |
| | | 63 | Maleate | 0.5 | 7.7 | 0.41 | -13.0 | 0.3 |
| | | 63 | Methylmalnote | 1 | 28.0 | 1.90 | -18.1 | 2.4 |
| | | 63 | Itaconate | 0.3 | 9.6 | 0.90 | -15.4 | 0.8 |
| | H8 | 91 | Uracil | 0.3 | 4.6 | 0.33 | -12.4 | 0.5 |
| | | 91 | Uridine | 0.3 | 3.9 | 0.53 | -10.8 | 0.7 |
| | | 91 | 5-Fluorouracil | 0.3 | 6.9 | 1.54 | -17.3 | 2.3 |
| | | 81 | Acetate | 2 | 103.0 | 4.72 | -14.3 | 1.7 |
| | | 81 | Butyrate | 2 | 133.0 | 13.10 | -13.5 | 4.0 |
| | | 81 | Propionate | 2 | 47.1 | 4.30 | -10.5 | 1.0 |
| | A4 | 87 | Uracil | 0.12 | 1.9 | 0.28 | -25.8 | 2.9 |
| | | 87 | Acetate | 0.5 | 50.3 | 5.76 | -8.7 | 0.7 |
| | | 87 | Butyrate | 0.5 | 29.9 | 2.36 | -17.2 | 1.2 |
| | | 72 | Propionate | 0.5 | 13.4 | 0.59 | -12.2 | 0.2 |
| | M8 | 70 | Uracil | 0.1 | 2.6 | 0.70 | -39.3 | 12.6 |
| | | 72 | Butyrate | 2 | 316.0 | 56.90 | -31.3 | 80.6 |
| | | 70 | Propionate | 8 | 633.0 | 104.00 | -0.9 | 0.2 |
| | | 70 | Acetate | 5 | N/A | N/A | N/A | N/A |

| | | | | | | | | |
|---|---|---|---|---|---|---|---|---|
| | M8-dm | 78 | Uracil | 0.12 | 2.6 | 0.32 | -23.6 | 0.9 |
| | | 78 | Butyrate | 2 | N/A | N/A | N/A | N/A |
| | | 78 | Propionate | 8 | N/A | N/A | N/A | N/A |
| | Apo-A4 | 68 | Uracil | 0.7 | 98.4x$10^{-3}$ | 6.6x$10^{-3}$ | -12.8 | 0.05 |
| | | 67 | Acetate | 3 | 74.3 | 5.92 | -9.9 | 1.14 |
| | | 67 | Butyrate | 3 | 104.0 | 8.81 | -5.5 | 0.57 |
| | | 68 | Propionate | 3 | 16.2 | 0.47 | -10.6 | 0.17 |
| | | 67 | Epinephrine | 3 | N/A | N/A | N/A | N/A |
| | Apo-A4_W160A | 68 | Uracil | 3 | 106.0 | 13.00 | -10.1 | 1.23 |
| | | 68 | Acetate | 3 | 87.7 | 6.41 | -6.8 | 0.51 |
| | Apo-A4_D205A | 66 | Uracil | 0.7 | N/A | N/A | N/A | N/A |
| | | 66 | Acetate | 3 | 51.4 | 5.36 | -4.6 | 0.31 |
| | Apo-A4_Y225A | 68 | Uracil | 0.7 | 54.8x$10^{-3}$ | 8.9x$10^{-3}$ | -13.5 | 0.09 |
| | | 68 | Acetate | 3 | N/A | N/A | N/A | N/A |
| | Apo-A4_Y273A | 68 | Uracil | 0.7 | 99.3x$10^{-3}$ | 23.4x$10^{-3}$ | -6.2 | 0.09 |
| | | 68 | Acetate | 3 | N/A | N/A | N/A | N/A |
| | Apo-A4_R116A, T145A, N178A, N180G | 67 | Uracil | 0.7 | N/A | N/A | N/A | N/A |
| | | 67 | Acetate | 3 | 87.1 | 4.15 | -11.5 | 0.83 |
| | | 67 | Epinephrine | 3 | 169.0 | 24.50 | -6.1 | 1.05 |
| **Competitive ITC** | Apo-A4 with uracil | 69 | Acetate | 3 | 63.7 | 4.85 | -12.0 | 1.18 |
| | Apo-A4 with acetate | 66 | Uracil | 0.7 | 75.1x$10^{-3}$ | 4.0x$10^{-3}$ | -14.0 | 0.04 |

**Supplementary Table 6. Crystallographic data collection and refinement statistics of A4 protein (PDB ID: 9HVJ).**

| | |
|---|---|
| **Data collection** | |
| Wavelength (Å) | 0.87313 |
| Resolution range | 46.54 - 1.464 (1.516 - 1.464) |
| Space group | $P\ 1\ 2_1\ 1$ |
| Unit cell | |
| a, b, c (Å) | 35.00, 93.08, 44.38 |
| α, β, γ (°) | 90.0, 92.2, 90.0 |
| Number of reflections | |
| Total | 330,830 (30,918) |
| Unique | 48,349 (4,659) |
| Multiplicity | 6.8 (6.6) |
| Completeness (%) | 99.07 (95.55) |
| Mean I/sigma(I) | 13.29 (3.31) |
| Wilson B-factor | 10.32 |
| R-merge | 0.1673 (0.9419) |
| R-meas | 0.1812 (1.023) |
| R-pim | 0.06886 (0.3939) |
| CC1/2 | 0.996 (0.732) |
| CC* | 0.999 (0.919) |
| **Refinement** | |
| Number of used reflections | |
| Total | 48,332 (4,659) |
| Free set | 2,416 (233) |
| R-work | 0.1681 (0.2338) |
| R-free | 0.1890 (0.2699) |
| Number of atoms | 2,579 |
| Macromolecules | 2,298 |
| Ligands | 16 |
| Solvent | 265 |
| Average B-factor | 15.83 |
| Macromolecules | 14.71 |
| Ligands | 11.18 |
| Solvent | 25.82 |
| R.M.S. deviations | |
| Bonds lengths (Å) | 0.007 |
| Bond angles (°) | 0.90 |
| Ramachandran | |
| Favored (%) | 97.85 |
| Allowed (%) | 2.15 |
| Outliers (%) | 0.00 |

Statistics for the highest-resolution shell are shown in parentheses.

### References


1. Broer, S. Intestinal amino acid transport and metabolic health. *Annu. Rev. Nutr.* **43**, 73-99 (2023).
2. Smith, E.A. & Macfarlane, G.T. Studies on amine production in the human colon: Enumeration of amine forming bacteria and physiological effects of carbohydrate and pH. *Anaerobe* **2**, 285-297 (1996).
3. Mager, L.F., *et al.* Microbiome-derived inosine modulates response to checkpoint inhibitor immunotherapy. *Science* **369**, 1481-1489 (2020).
4. Meier, K.H.U., *et al.* Metabolic landscape of the male mouse gut identifies different niches determined by microbial activities. *Nat. Metab.* **5**, 968-980 (2023).
5. Psarra, T.A., Batzias, G.C., Peeters, T.L. & Koutsoviti-Papadopoulou, M. The gastrointestinal effects that may follow the administration of theophylline reflect the pharmacodynamic profiles of both the parent drug and its metabolites. *Fundam. Clin. Pharmacol.* **24**, 171-180 (2010).
6. Connors, J., Dawe, N. & Van Limbergen, J. The role of succinate in the regulation of intestinal inflammation. *Nutrients* **11**(2018).
7. Hughes, E.R., *et al.* Microbial respiration and formate oxidation as metabolic signatures of inflammation-associated dysbiosis. *Cell Host Microbe* **21**, 208-219 (2017).
8. Kircher, B., *et al.* Predicting butyrate- and propionate-forming bacteria of gut microbiota from sequencing data. *Gut Microbes* **14**, 2149019 (2022).
9. Gargari, G., Taverniti, V., Koirala, R., Gardana, C. & Guglielmetti, S. Impact of a multistrain probiotic formulation with high bifidobacterial content on the fecal bacterial community and short-chain fatty acid levels of healthy adults. *Microorganisms* **8**(2020).
10. Xie, J., *et al.* Fructose metabolism and its role in pig production: A mini-review. *Front. Nutr.* **9**, 922051 (2022).
11. Yang, J., *et al.* Biphasic chemotaxis of *Escherichia coli* to the microbiota metabolite indole. *Proc. Natl Acad. Sci. USA* **117**, 6114-6120 (2020).
12. Anton, B.P., Fomenkov, A., Raleigh, E.A. & Berkmen, M. Complete genome sequence of the engineered *Escherichia coli* SHuffle strains and their wild-type parents. *Genome Announc.* **4**(2016).
13. Woodcock, D.M., *et al.* Quantitative evaluation of *Escherichia coli* host strains for tolerance to cytosine methylation in plasmid and phage recombinants. *Nucleic Acids Res.* **17**, 3469-3478 (1989).
14. Ames, P., Studdert, C.A., Reiser, R.H. & Parkinson, J.S. Collaborative signaling by mixed chemoreceptor teams in *Escherichia coli*. *Proc. Natl Acad. Sci. USA* **99**, 7060-7065 (2002).
15. Sourjik, V. & Berg, H.C. Functional interactions between receptors in bacterial chemotaxis. *Nature* **428**, 437-441 (2004).
16. Burón-Barral, M.D.C., Gosink, K.K. & Parkinson, J.S. Loss- and gain-of-function mutations in the F1-HAMP region of the *Escherichia coli* aerotaxis transducer Aer. *J. Bacteriol.* **188**, 3477-3486 (2006).
17. Bi, S., Jin, F. & Sourjik, V. Inverted signaling by bacterial chemotaxis receptors. *Nat. Commun.* **9**, 2927 (2018).
